# Ablation of polysaccharide breakdown in *Bacteroides thetaiotaomicron* prevents cross-feeding and growth of *Salmonella* Typhimurium in the mouse gut

**DOI:** 10.1101/2025.08.08.669261

**Authors:** Lena Amend, Anja Stock, Marie Wende, Lisa Osbelt-Block, Gesa Martens, Meina Neumann-Schaal, Till Strowig, Alexander J. Westermann

## Abstract

Pathogens invading the intestine compete for nutrients with the resident microbiota. However, there is evidence that commensal members of the gut also provide nutritional resources to enteropathogens and thus promote their outgrowth. In this study, we investigated metabolic cross-feeding mechanisms between the abundant gut commensal *Bacteroides thetaiotaomicron* and the model enteropathogen *Salmonella enterica* serovar Typhimurium. We discovered that the processing of various dietary and host-derived glycans by *B. thetaiotaomicron* liberated building blocks available to *Salmonella* and identified a range of cross-fed metabolites. Interfering with polysaccharide degradation in *B. thetaiotaomicron* by genetic manipulation of specific polysaccharide utilization loci (PUL) inhibited pathogenic cross-feeding, both *in vitro* and in a gnotobiotic mouse model. Our findings highlight the complex metabolic commensal-pathogen interaction in the intestine and propose the disruption of polysaccharide breakdown as a potential microbiota-centric strategy to intervene in intestinal infections.

## INTRODUCTION

*Salmonella enterica* is an important enteric pathogen infecting humans and livestock (Eng *et al*., 2015; Han *et al*., 2024). Within the mammalian gastrointestinal (GI) tract, *Salmonella*’s metabolic versatility as a nutrient generalist is crucial for its survival and ability to cause disease (Eisenreich *et al*., 2010; Rivera-Chávez and Bäumler, 2015). The pathogen entertains multiple, partially redundant pathways to catabolize simple sugars, and inhibition of individual metabolic enzymes is typically insufficient to decrease *Salmonella* virulence (Becker *et al*., 2006). Therefore, rather than interfering with *Salmonella* metabolic activities, can one restrict the carbohydrate substrates that feed into these pathways in order to counteract *Salmonella* infection?

The metabolic environment in the mammalian gut is fundamentally shaped by the intestinal microbiota. By deploying carbohydrate-active enzymes (CAZymes), commensal gut bacteria can catabolize complex dietary and host-derived glycans (Wardman *et al*., 2022). Members of *Bacteroides*—a dominant genus in the mammalian intestine (Arumugam *et al*., 2011)—are equipped with a multitude of genes for different CAZymes (Kaoutari *et al*., 2013; Lapébie *et al*., 2019). Typically, these CAZymes are genetically linked to the cognate transporters and regulatory proteins in multi-gene operons referred to as polysaccharide utilization loci (PULs) (Bjursell, Martens and Gordon, 2006). For example, *Bacteroides thetaiotaomicron* (*B. theta*) encodes 96 distinct PULs (cazy.org, (Drula *et al*., 2021)), endowing this microbiota model organism with the metabolic flexibility needed to adapt to carbon source fluctuations in the intestinal environment (Wexler, 2007; Wexler and Goodman, 2017; Porter, Luis and Martens, 2018).

The molecular mechanism of polysaccharide processing and import by PUL systems was first resolved for the starch-catabolic PUL66 of *B. theta* (Anderson and Salyers, 1989a, 1989b). This “starch-utilization system” (Sus) encompasses eight proteins, including the regulatory and sensory protein SusR, the outer membrane proteins SusC-SusG that mediate binding (SusDEF), hydrolysis (SusG), and import (SusC) of the substrate into the periplasm, as well as periplasmic enzymes (SusAB) for further hydrolytic breakdown into maltose and glucose—the monosaccharide constituents of starch (Foley, Cockburn and Koropatkin, 2016).

While there is a debate as to whether glycan degradation by *Bacteroides* spp. is primarily selfish or deliberately co-operative (Culp and Goodman, 2023), an increasing number of examples suggest synergistic cross-feeding interactions within the intestinal microbiota. The main breakdown products of fibres that are liberated by the activity of primary degraders, such as *B*. *theta,* are short-chain fatty acids (SCFAs) (Cummings *et al*., 1987), particularly acetate and propionate (Louis and Flint, 2017; Frolova *et al*., 2022). SCFAs play a beneficial role in maintaining intestinal barrier integrity and systemic health, e.g., by exerting anti-inflammatory or immune-regulatory activities (Mukhopadhya and Louis, 2025), but may also be consumed by other intestinal bacteria. Similarly, *Bacteroides*-derived fermentation products such as succinate and formate (Louis and Flint, 2017), as well as oligo– and monosaccharides, (Rakoff-Nahoum, Coyne and Comstock, 2014) can be consumed by members of the intestinal community (Payling *et al*., 2020).

There is also evidence that gut microbiota-derived metabolites can foster the outgrowth of invading pathogens. For example, the intestinal pathobiont *Clostridioides difficile* and pathogenic *Escherichia coli* thrive on sialic acid released through the breakdown of host mucus by *Bacteroides* spp. (Ng *et al*., 2013; Huang *et al*., 2015), and *B. theta*-derived succinate promotes the proliferation of *C. difficile* (Ferreyra *et al*., 2014). Likewise, the model enteric pathogen *Salmonella enterica* serovar Typhimurium (*S.* Tm) exploits several microbiota-derived metabolites: it expands on *B. theta*-derived sialic acid and fucose (Ng *et al*., 2013), propionate (Shelton *et al*., 2022), and the SCFA intermediates succinate (Spiga *et al*., 2017) and 1,2-propanediol (Faber *et al*., 2017). This supports the potential for rational intervention of catabolic processes in the indigenous microbiota as an anti-infective strategy to prevent or treat intestinal diseases. However, such efforts are currently hindered by our limited understanding of the molecular basis of commensal-pathogen cross-feeding interactions in the gut.

In the present study, we systematically investigate cross-feeding mechanisms between the two model organisms, *B. theta* and *S.* Tm, across a comprehensive panel of dietary and host-derived polysaccharides. Next to dissecting the respective polysaccharide degradation machineries of *B. theta*, we identify *B. theta* pre-processed polysaccharides cross-fed to, and exploited by *S*. Tm. Disrupting glycan cross-feeding from *B. theta* to *S.* Tm by genetic inactivation of specific PUL genes prevents *S*. Tm outgrowth in a gnotobiotic mouse model. Together, this study establishes causality of glycan cross-feeding for bacterial virulence.

## RESULTS

### PUL expression profiling during B. theta growth on a variety of complex polysaccharides

*B. theta* is a model organism for primary degraders in the human intestine and is capable of breaking down dietary fibers that are indigestible to the host and most companion microbes (Martens, Chiang and Gordon, 2008; Martens *et al*., 2011). Here, we grew *B. theta* type-strain VPI-5482 in VB minimal medium (Varel and Bryant, 1974) supplemented with each one out of a panel of 30 distinct microbiota-accessible carbohydrates (MACs), clustered into seven polysaccharide classes (Suppl. Table 1), as the sole carbon source. In line with previous data (Martens, Chiang and Gordon, 2008; Martens *et al*., 2011), *B. theta* grew on multiple polysaccharides, including all tested glycans from the groups of starches, fructans, pectins, as well as bacterial and fungal cells (Figure 1A & S1A-G). In contrast, *B. theta* grew on only one (namely xylan, derived from corncob) out of nine tested hemicelluloses and failed to grow on the algal glucan laminarin and host-derived heparin.

**Figure 1:**
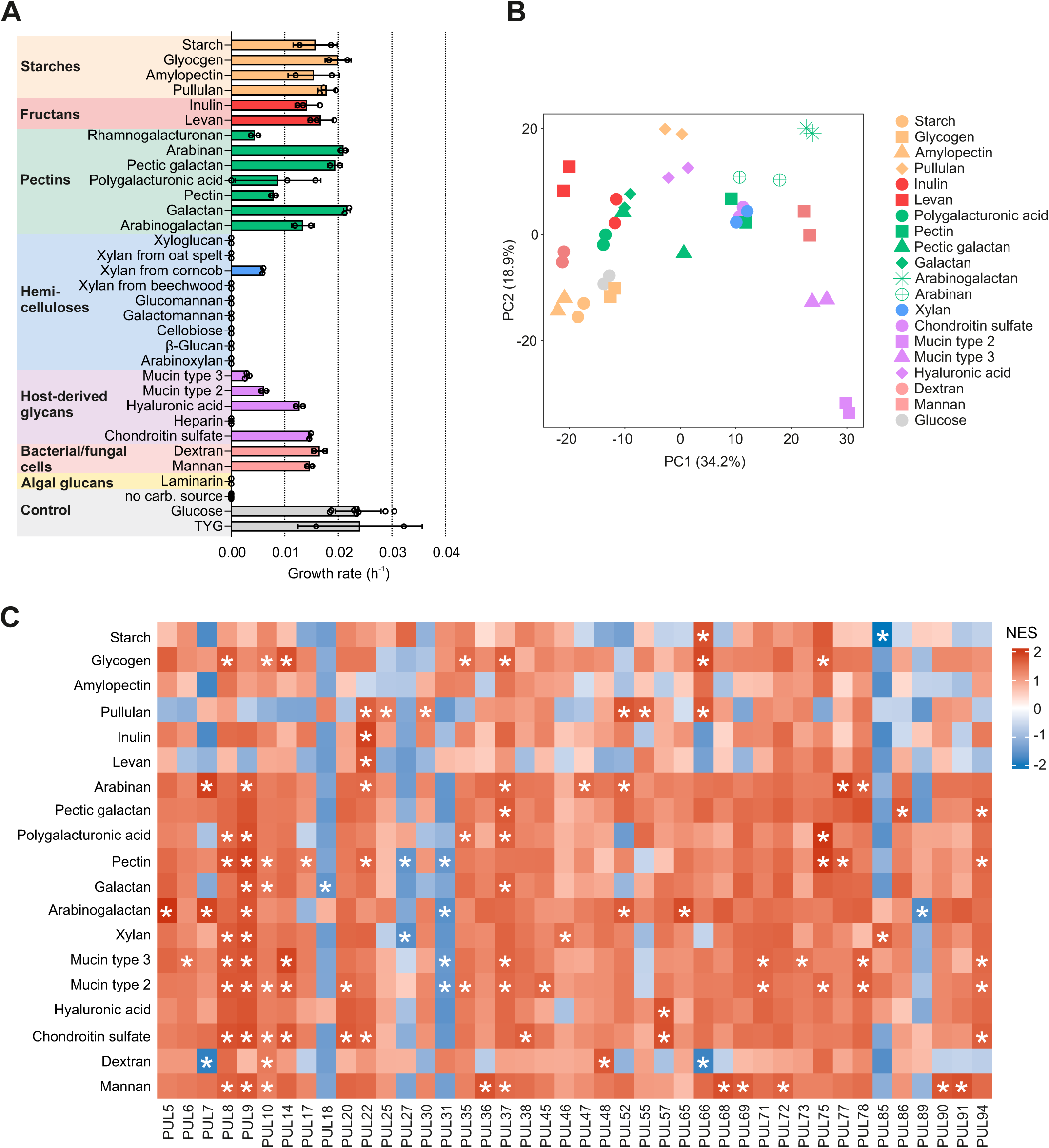
*B. theta* growth and PUL expression on a variety of polysaccharides. A: Anaerobic growth rates of *B. theta* grown in minimal VB medium supplemented with different diet-or host-derived polysaccharides. The mean and standard deviation from three biological replicates are plotted as bars and error bars, respectively, with each replicate depicted as a single dot. B: Global gene expression of *B. theta* grown in VB medium supplemented with the 19 indicated polysaccharides and glucose as control. Shown is a PCoA plot with each data point representing one biological replicate. Chemically related carbohydrates are depicted in the same color, while represented by different symbols. C: Gene set enrichment analysis of the 96 literature-derived PULs annotated using the CAZy PULDB (cazy.org). Lower levels of PUL expression in *B. theta* grown in polysaccharides compared to glucose are represented in shades of blue and higher expression in red. NES: normalized enrichment score, FDR: false discovery rate. An asterisk (*) indicates an FDR < 0.05.

We measured global gene expression of *B. theta* grown on each of the 19 polysaccharides that supported growth. To this end, total RNA was extracted from *B. theta* cultures grown to early stationary phase (sampling ODs are listed in Suppl. Table 2), depleted from ribosomal RNA, converted into cDNA libraries, and sequenced. A *B. theta* culture grown in glucose-containing medium served as a reference control. Principal component analysis (PCoA) of the obtained data revealed that the transcriptomes of biological replicates clustered (Figure 1B), ruling out any major batch effects. Moreover, *B. theta* grown on galactan, polygalacturonic acid, and pectic galactan, on arabinogalactan and arabinan, or on starch, glycogen, and amylopectin formed specific clusters, suggesting that *B. theta* employs overlapping gene sets to grow on chemically related substrates. *B. theta* samples grown on porcine gastric mucins (crude [type 2] mucin or a partially purified variant thereof [type 3]) segregated from the other polysaccharides, reflecting their unique glycan structure.

Differential gene expression and gene set enrichment analyses corroborate previously reported associations between specific PULs and their known substrates (Martens, Chiang and Gordon, 2008; Martens *et al*., 2011; Cuskin *et al*., 2015; Ndeh *et al*., 2017, 2025; Luis *et al*., 2018; Ryan *et al*., 2024). Generally, growth on complex heteropolysaccharide classes (composed of different sugar monomers), including pectins and host-derived glycans, was accompanied by the upregulation of multiple PUL systems (Figure 1C, Suppl. Figure S2), suggesting the concerted action of multiple PULs to enable the breakdown of complex carbohydrates. For example, 8 or 12 distinct PULs were significantly (FDR<0.05) upregulated during growth on pectin or type 2 mucin, respectively. When *B. theta* grew on α-glucans (i.e., glycans entirely composed of glucose monomers linked by α-glycosidic bonds), including starch, amylopectin, glycogen, pullulan, and dextran, as well as the fructans inulin and levan (that both consist exclusively of fructose monomers), it induced only a small subset of PULs, while downregulating others. Growth on inulin and levan evoked the induction of only PUL22, hyaluronic acid the specific upregulation of PUL57, and starch the strong upregulation of only PUL66. Conversely, our analysis revealed that PUL18 and PUL31 were predominantly downregulated in the presence of any complex polysaccharide compared to glucose. Both PULs are relatively short, comprising three (PUL18; namely *BT1551*: hypothetical protein, *BT1552*: SusC, *BT1553*: SusD) or four genes (PUL31; *BT2362*: SusC, *BT2363*: SusD, *BT2364*: SusC, *BT2365*: SusD), respectively, while lacking genes for regulatory elements or CAZymes. In line with previous findings (Liu *et al*., 2021), this suggests that some PULs of *B. theta* are responsible for the transport/uptake of simple sugars that derive from the partial breakdown of polysaccharides by unrelated PULs.

Overall, this comprehensive transcriptome dataset illustrates polysaccharide-specific gene expression patterns in *B. theta*, advancing our understanding of polysaccharide metabolism and the corresponding degradation machinery in *B. theta*. The RNA-seq data are publicly available through our previously launched server ‘Theta-Base’ (Ryan *et al*., 2020, 2024) and will serve as a valuable resource for the microbiome research community.

### Salmonella fails to grow on complex polysaccharides, unless they are pre-processed by B. theta

*S.* Tm is a facultative anaerobic, food-borne gut pathogen belonging to the *Enterobacteriaceae*. As expected, *S.* Tm strain SL1344 was unable to grow on any of the tested polysaccharides—both under aerobic and anaerobic conditions—whereas we observed robust growth in control media (rich Luria-Bertani [LB] medium, M9 supplemented with glucose, or M13 minimal medium) (Figure 2A, B, Suppl. Figure S3A-G). To test whether polysaccharide pre-processing by *B. theta* would permit growth of *S.* Tm, a spent media assay was devised. *B. theta* was grown in M13 medium supplemented with single polysaccharides until reaching stationary phase (sampling ODs are listed in Suppl. Table 2), the supernatant was harvested, filter-sterilized, and inoculated with *S.* Tm, and bacterial growth was monitored (Figure 2B). The pre-processing of all 19 tested polysaccharides by *B. theta* enabled aerobic growth of *S.* Tm (Figure 2B, Suppl. Figure S4 A-G). In the analogous assay under anaerobic conditions, *S*. Tm outgrowth was not observed, probably due to increased nutrient depletion or the accumulation of inhibitory substances produced by *B. theta* during the stationary growth phase. This is because the simultaneous co-cultivation of both bacterial species in M13 supplemented with the same polysaccharides in co-culture plates did result in *S*. Tm growth under anaerobic conditions (Figure 2C, Suppl. Figure S5A-S). We conclude that cross-feeding of *S*. Tm by *B. theta* allows the outgrowth of this pathogen in the presence of oxygen and to a lesser extent within an anaerobic environment that resembles the physiological conditions in the intestine.

**Figure 2:**
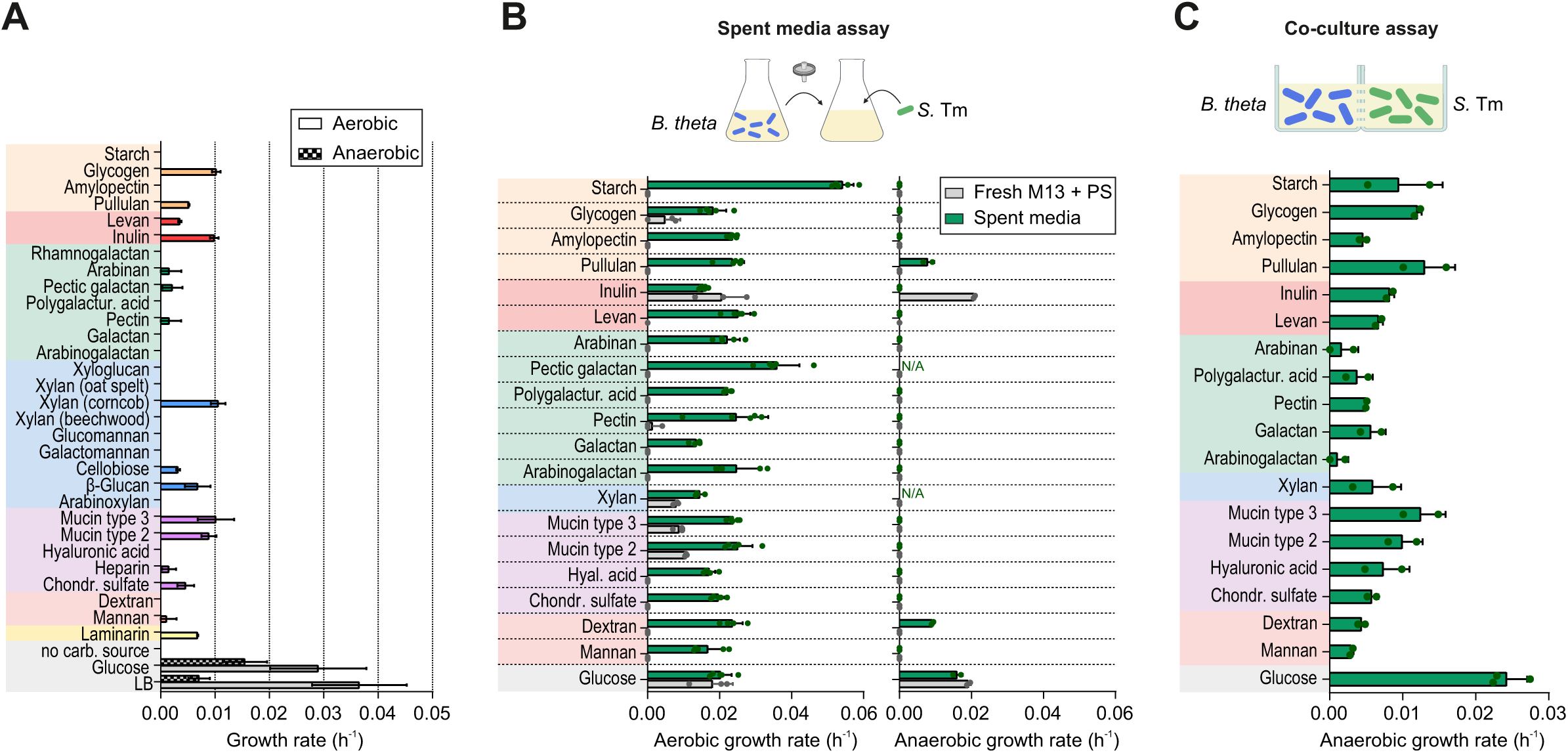
*S.* Tm fails to grow on polysaccharides, but can utilize metabolites liberated by *B. theta*. A: Aerobic and anaerobic growth rates of *S.* Tm grown in minimal M9 medium supplemented with the same panel of polysaccharides used in Figure 1. The mean and standard deviation from two biological replicates are plotted as bars and error bars, with each replicate depicted by a single dot. Note that the commercial inulin, corncob xylan, and mucin type 2 or 3 powders that were used in this assay contained free residues of various simple sugars or amino acids (see metabolomics data in Suppl. Figure 6), suggesting that the subtle *S.* Tm growth observed here was probably supported by these less complex molecules rather than the respective glycan. B: Spent media assay. Upper: experimental scheme. *B. theta* is grown to stationary phase in minimal M13 medium supplemented with single polysaccharides. Supernatants are filter-sterilized and inoculated with *S.* Tm. *S*. Tm density is monitored after 24 h of growth under aerobic or anaerobic conditions. Lower: mean *S*. Tm growth rates. Aerobic growth rate determined from 5 biological replicates in spent media (identical samples as used in Figure 3 for metabolome analysis) and from 3 biological replicates in fresh M13; anaerobic growth determined from 2 biological replicates. Each replicate is represented by a single dot. Error bars denote standard deviation from the mean. C: Co-culture assay. Upper: experimental scheme. *B. theta* and *S.* Tm are inoculated into M13 medium supplemented with single polysaccharides and dispensed into each two wells separated by a semi-permeable membrane (pore size = 0.2 µm). Growth is monitored simultaneously for *B. theta* and *S*. Tm (growth curves in Suppl. Figure 6). Lower: anaerobic growth rate of *S.* Tm co-cultured with *B. theta* under the indicated conditions. Bars denote the mean and error bars the standard deviation of each two biological replicates (represented by individual dots).

In the absence of oxygen, *S.* Tm relies on alternative electron acceptors to perform anaerobic respiration (Gennis, Stewart and Neidhardt, 1996). Nitrate, which is increasingly secreted by the host upon intestinal infection, can serve as alternative electron acceptor for *S.* Tm (Barrett and Riggs, 1982). To assess a potential effect of nitrate on *S.* Tm growth in an anaerobic environment, the spent media assay was repeated in the additional presence of 40 mM sodium nitrate (Winter *et al*., 2013; Faber *et al*., 2017; Shelton *et al*., 2022). Nitrate addition diminished *S*. Tm growth in fresh M13 supplemented with glucose and inulin, but enabled slight *S*. Tm growth in fresh M13 containing glycogen or mucin and several *B. theta*-processed polysaccharides (Suppl. Figure 4H). This implies that the access to alternative electron acceptors can enhance anaerobic growth of *S*. Tm on specific oligosaccharides. Altogether, these data demonstrate that *B. theta* pre-degradation of MACs opens new nutrient sources for *S.* Tm.

### Identification of metabolites released by B. theta as a result of glycan consumption that are consumed by S. Tm

To pinpoint the metabolites that are produced by *B. theta*—and hence become available to *S*. Tm— during polysaccharide degradation, a targeted metabolite analysis was conducted. Spent media from *B. theta* grown in M13 supplemented with each one of the 18 different polysaccharides, as well as glucose, was harvested before and after incubation with *S.* Tm for 24 h (Figure 2B). We measured the levels of 77 metabolites, including select carbohydrates, amino acids, branched short chain fatty acids (BSCFAs), SCFAs and their respective intermediates using GC/LC-MS (raw data in Suppl. Table 3). The measurement of blank M13 with the individual carbon sources as a quality control detected various amino acids in the majority of the commercial polysaccharide charges, especially so in the case of the four host-derived glycans (chondroitin sulfate, hyaluronic acid, mucin types 2 and 3) (Suppl. Figure S6). Furthermore, the inulin and xylan batches inherently contained high levels of their respective mono– and disaccharide constituents (namely fructose and xylobiose) (Suppl. Figure S6), and were thus precluded from further analyses.

Global metabolic profiles of *B*. *theta* grown on the different glycans were visualized by PCoA. Overall, the metabolomes of *B*. *theta* grown in M13 supplemented with mucins segregated from those with the other polysaccharides (Figure 3A). Metabolite class-specific PCoA (for amino acids, volatiles, or non-volatiles) indicated that the segregation between mucins and the remaining glycans was mostly driven by a differential production of amino acids (Suppl. Figure 7A) and volatile compounds (Suppl. Figure 7B), without major differences with respect to non-volatile (predominantly saccharide) profiles (Suppl. Figure 7C). Excluding the mucin samples from PCoA revealed the tight clustering of biological replicates and of structurally related glycans (starch, amylopectin, dextran; pectin and polygalacturonic acid; [pectic] galactan) (Figure 3B), reflecting the pattern observed for the corresponding transcriptomes (Figure 1).

**Figure 3:**
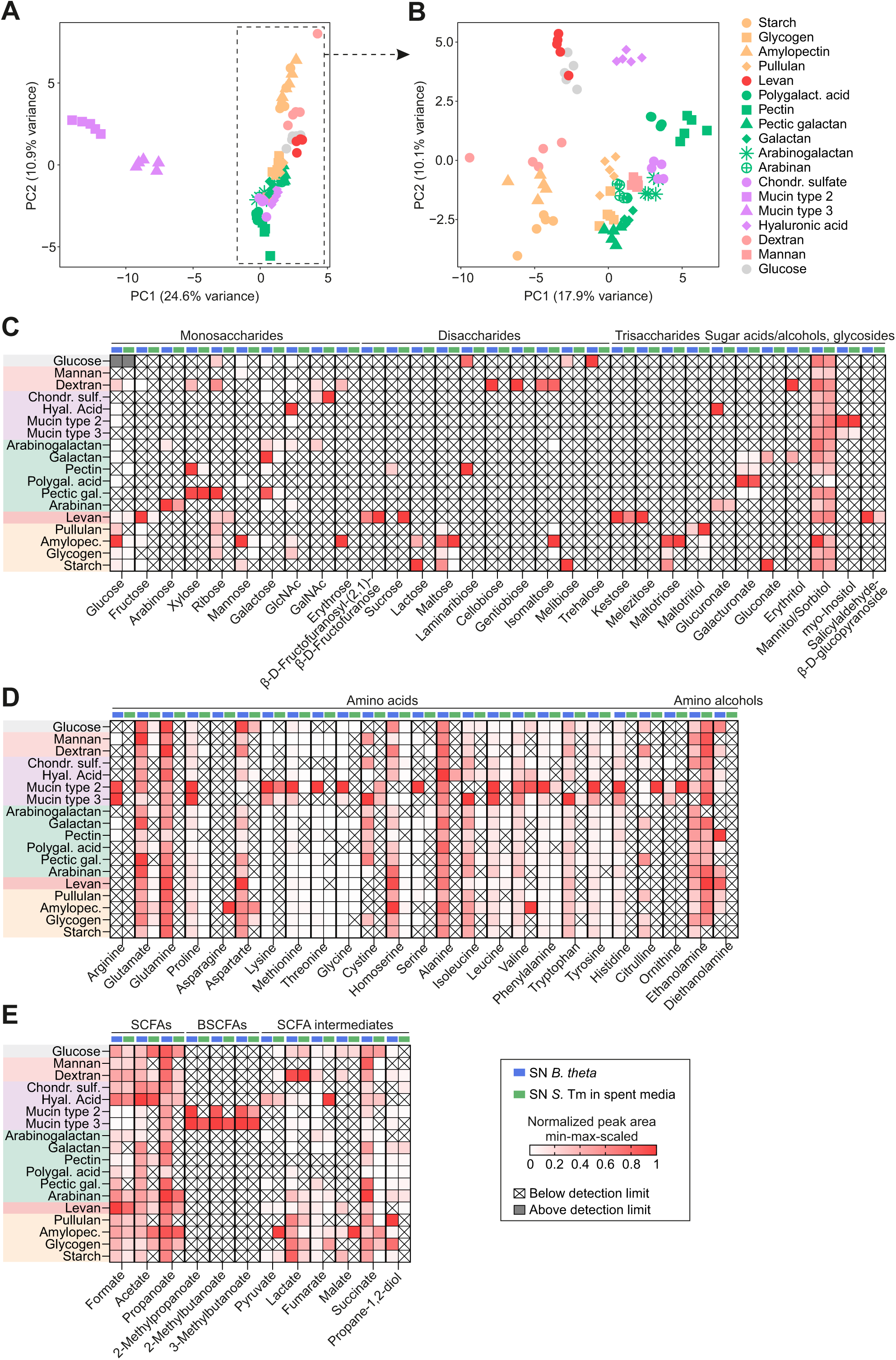
Metabolites generated or consumed by *B. theta* or *S*. Tm, respectively. A-B: PCoA plot of global metabolic profiles of *B. theta* grown in M13 supplemented with different polysaccharides or glucose as control, either including (A) or excluding (B) the data derived from mucin cultures. Each data point represents one biological replicate (out of 5). Polysaccharide subclasses grouped by color. C-E: Detection of mono-, di-, and trisaccharides, sugar acids, sugar alcohols, and glycosides (C), of amino acids and amino alcohols (D), and of SCFAs, BCFAs, and SCFA intermediates (E) in the supernatants of *B. theta* grown in M13 supplemented with 17 different glycans or glucose (blue) and the respective samples after 24 h of *S.* Tm growth (green). Metabolome data of xylan– and inulin-derived samples were excluded due to high concentrations of monosaccharides in the corresponding blank samples. Data represent min-max-scaled peak areas (or, in case of separately measured formate, concentration in g/L) averaged over the 5 biological replicates. White crossed-out cells indicate that a metabolite was below the detection limit and grey cells, that it exceeded the detection limit.

Out of the 77 analyzed metabolites, 68 were detected in at least one supernatant fraction (Figure 3C-E). The accumulation of carbohydrates in the *B. theta* culture supernatants was highly specific and driven by the respective glycan substrate (Figure 3C). For instance, growth of *B. theta* in M13 supplemented with galactan resulted in the accumulation of glucose, galactose, galacturonate, gluconate, and erythritol; growth on arabinan released free arabinose, galactose, GlcNAc, and glucuronate; growth on hyaluronic acid led to the production of glucose, GlcNAc, and glucuronate. *B. theta* consumption of mucins released only a few different saccharides, and so in low concentrations. Following growth of *S.* Tm for 24 hours, the vast majority of *B. theta*-liberated carbohydrates was depleted from the cultures (Figure 3C). Only in a few cases did *S.* Tm not use up the carbohydrates released by *B. theta*. For example, isomaltose was retained in *B. theta*-spent media derived from dextran, myo-inositol was not depleted in spent media derived from mucins, galacturonate in spent media from galactan, pectin, polygalacturonic acid, and pectic galactan, and inulobiose (β-D-Fructofuranosyl-[2,1]-β-D-Fructofuranose) and kestose were retained in supernatants from levan. Similarly, a sugar alcohol (mannitol or sorbitol), which was nonspecifically produced by *B. theta* across all glycan substrates, was not consumed by *S.* Tm.

Contrasting the specific accumulation of carbohydrates in the *B. theta* supernatants, amino acids accumulated under all conditions and their levels peaked when *B. theta* was grown on both types of mucins, probably due to the polypeptide backbone of these glycoproteins (Suppl. Figure 6). All quantified amino acids and the amino alcohol diethanolamine were consistently depleted following incubation with *S.* Tm (Figure 3D). In contrast, ethanolamine increased following *S.* Tm incubation, suggesting this particular amino alcohol to not be consumed, but rather produced by *S.* Tm.

The BSCFAs 2-methylpropanoate and 2-methylbutanoate were only released by *B. theta* during growth on mucins, presumably as a result of the fermentation of branched chain amino acids (Smith and Macfarlane, 1998; Aguirre *et al*., 2016) contained in the input mucin mixtures (Suppl. Figure S6). The SCFAs formate, acetate, and propanoate were released by *B. theta* in presence of all tested polysaccharides (Figure 3E). The consumption of *B. theta*-derived SCFAs, BSCFAs, and SCFA intermediates by *S.* Tm mainly depended on the polysaccharide source (Figure 3E). For instance, *S.* Tm depleted propanoate from the spent media derived from mannan, mucins, pectin, pullulan, and starch. In contrast, propanoate from *B. theta* grown on glucose, hyaluronic acid, arabinan, levan, and amylopectin was barely utilized by *S.* Tm.

Taken together, the metabolomic data support the extensive carbon cross-feeding between *B. theta* and *S.* Tm. In particular, *B*. *theta*-liberated mono– and disaccharides are preferably consumed by *S*. Tm.

### Genetic inactivation of glucose import in S. Tm is insufficient to prevent pathogen outgrowth in a nutrient-rich environment

Having uncovered that simple sugars are readily cross-fed from *B. theta* to *S*. Tm, we next asked whether inhibition of carbohydrate uptake would be sufficient to disrupt the growth of this pathogen under these conditions. The simultaneous deletion of two different glucose-importing phosphotransferase systems (PtsG, ManXYZ) and of glucokinase (Glk; catalyzes the first step in glucose catabolism) in *S*. Tm (Fig. 4A) effectively inhibited the outgrowth of the resulting mutant in minimal medium containing glucose as the sole carbon source (Fig. 4B), as previously reported (Götz and Goebelt, 2010). In stark contrast, growth of the same triple mutant was not compromised in spent media from *B. theta* pre-grown in the same minimal glucose medium (Fig. 4C) and only subtly delayed in spent medium from *B. theta* cultures grown on dextran (Fig. 4D). Metabolomic measurements pointed at the expected increase in glucose levels in the spent medium after the outgrowth of ΔΔΔ*ptsG*/*manXYZ*/*glk* compared to wild-type *S*. Tm and a concomitant depletion of alternative carbon sources of *Salmonella*, particularly succinate (Spiga *et al*., 2017) and specific amino acids (including glutamate and aspartate; (Zampieri *et al*., 2019)) (Fig. 4E, F, Suppl. Table 4). These findings highlight the metabolic robustness of *S*. Tm, in line with the difficulty to intervene with *Salmonella* fitness by targeting select metabolic enzymes in this pathogen (Becker *et al*., 2006). In contrast, we next evaluated the potential of intervening at a preceding step, namely inactivating specific metabolic processes in *B. theta*, thereby possibly limiting the availability of *Salmonella*-accessible nutrients.

**Figure 4:**
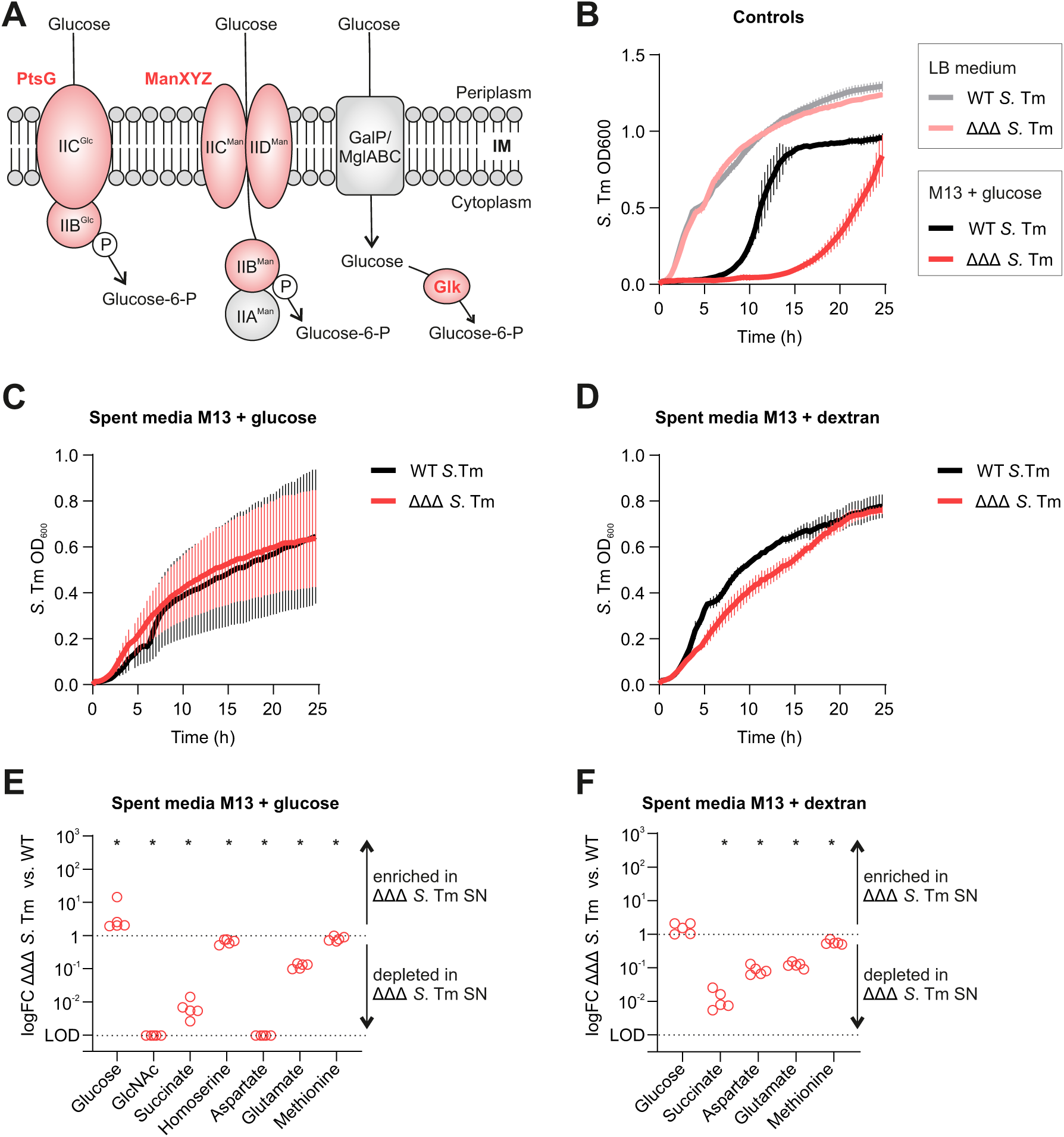
Glucose import is dispensable for *S*. Tm growth on *B*. *theta*-spent medium. A: Schematic of *Salmonella* monosaccharide uptake systems (in red) that are inactivated in the ΔΔΔ*ptsG*/*manXYZ*/*glk* mutant (dubbed ‘ΔΔΔ’). This previously described mutant combines the deletions of *ptsG* (encoding IIBC^Glc^ of the glucose-specific phosphotransferase system [PTS] transporter), *manXYZ* (encoding IIBCD^Man^ of the mannose PTS, which can also import glucose), and *glk*, the gene encoding the glucosekinase that phosphorylates non-PTS transported glucose in the cytosol, in the *S*. Tm 14028 strain background (Götz and Goebelt, 2010). ‘IM’ refers to inner membrane and ‘P’ to phosphate. Figure adapted from (Bowden *et al*., 2014). B: Growth kinetics of *S*. Tm wild-type and ΔΔΔ in rich (LB) or minimal glucose medium. Data refer to the mean and standard deviation from each 5 biological replicates. While we observed comparable growth patterns between ΔΔΔ and wild-type *S*. Tm in LB, the triple mutant displayed an extended lag phase during growth in M13 supplemented with the sole carbon source glucose. After ∼15 h, the triple mutant started to grow, presumably due to the accumulation of genomic mutations (Götz and Goebelt, 2010). (Note that, in line with Figure 2A, neither strain grows on minimal dextran-containing medium.) C, D: Growth kinetics of the same *S*. Tm strains in spent medium derived from *B*. *theta* cultures grown in M13 medium with either glucose (C) or dextran (D) as the sole carbon source. Data refer to the mean and standard deviation from each 5 biological replicates. E, F: Targeted metabolome analysis of the spent media assays shown in panels C and D. Endpoint quantification at 24 h of *S*. Tm growth in spent medium from M13 with either glucose (E) or dextran (F). The levels of the metabolites shown were reduced compared to their input concentration (time point 0 of the spent media assay) and, additionally, significantly (p ≤ 0.05 according to a paired, non-parametric Wilcoxon matched-pairs signed rank test; indicated by an asterisk [*]) altered between ΔΔΔ vs. wild-type *S*. Tm culture supernatants. Single dots reflect the pairwise logFC over 5 biological replicates. LOD, limit of detection.

### Genetic inactivation of B. theta PULs 22 and 48 disrupts S. Tm cross-feeding in vitro

Aiming to suppress the production of catabolites released from polysaccharide breakdown, we sought to predict *B. theta* PULs critical for *S*. Tm cross-feeding based on our collective datasets. Our transcriptomics data supported previous reports (Martens, Chiang and Gordon, 2008; Sonnenburg *et al*., 2010; Martens *et al*., 2011; Wong *et al*., 2023) that PUL22 is required for the breakdown of levan, PUL48 for dextran breakdown, PUL57 for hyaluronic acid breakdown, and PUL75 for the processing of polygalacturonic acid (Figure 1C, Suppl. Figure S2). Moreover, we identified multiple metabolites (glucose, fructose, mannose, GlcNAc, kestose, melezitose, salicylaldehyde-β-D-glucopyranoside, cellobiose, gentiobiose, erythritol, glucuronate, galacturonate) that specifically accumulated upon *B. theta* growth on these polysaccharides and were depleted upon incubation with *S.* Tm. In combination, this suggested PUL22, PUL48, PUL57, and PUL75 as promising target candidates to block *S.* Tm cross-feeding. To test this possibility, we created deletion mutants of the cognate PUL-encoded transcriptional activators: BT1754 (HTCS), BT3091 (SusR) (Figure 5A), BT3334 (HTCS), and BT4111 (HTCS) (Suppl. Figure S9A). Reassuringly, the regulator-deficient mutants were unable to grow on their respective substrate (Suppl. Figure S8A-D), while complementation in *trans* (Suppl. Figure S8F-I) restored their growth to wild-type levels.

**Figure 5:**
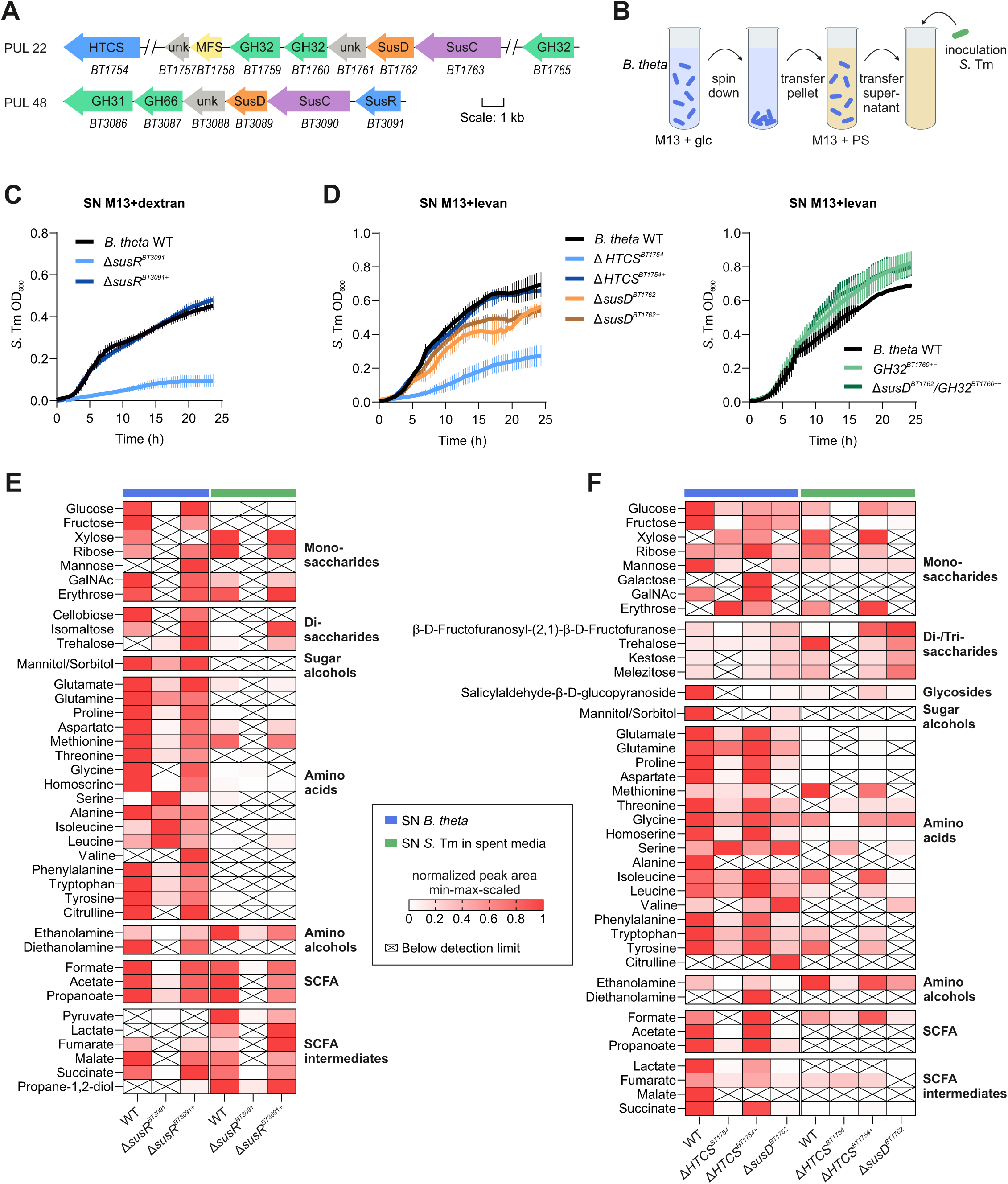
Genetic manipulation of PULs 22 and 48 affects cross-feeding to *S.* Tm. A: Genetic organization of levan-catabolic PUL22 and dextran-catabolic PUL48 in *B. theta*. Locus tags and, where applicable, annotated protein functions (from cazy.org) are indicated. HTCS: hybrid two component system, GH: glycoside hydrolase, MFS: major facilitator superfamily, unk: unknown (hypothetical protein). PUL organization was plotted using Gene Graphics (Harrison, Crécy-Lagard and Zallot, 2017). B: Experimental workflow: *B. theta* cells are grown anaerobically in M13 medium containing glucose. Bacterial pellets are collected after 24 h, transferred to M13 medium supplemented with either levan or dextran, and cultures are incubated therein for another 4 (levan) or 8 h (dextran). The resulting supernatant is filter-sterilized and inoculated with *S.* Tm, whose growth is then monitored under aerobic conditions. C: Growth curves of *S.* Tm in the supernatant from *B. theta* wild-type, Δ*susR^BT3091^* or the corresponding SusR-complemented strain (Δ*susR^BT3091+^*) that had been collected after 8 h of incubation in M13-dextran medium. The data refer to the mean and standard deviation from each 5 biological replicates. D: Growth curves of *S.* Tm in the supernatant from *B. theta* wild-type, Δ*HTCS^BT1754^,* Δ*susD^BT1762^* or the respective trans-complemented strains (left) that had been collected after 4 h of incubation in M13-levan medium, as well as in those from *GH32^BT1760++^* and Δ*susD^BT176^*/*GH32^BT1760^*^++^ overexpression strains (right) collected after 8 h of incubation in M13-levan medium. The data refer to the mean and standard deviation from either 5 (left) or 3 (right) biological replicates. E: Metabolites measured in the supernatants of *B. theta* wild-type, Δ*susR^BT3091^* or Δ*susR^BT3091+^* strains grown in dextran, prior to (blue) or after (green) the incubation with *S*. Tm for 24 h. F: Metabolites measured in the supernatants of *B. theta* wild-type, Δ*HTCS^BT1754^*, Δ*HTCS^BT1754+^*, and Δ*susD^BT1762^* grown in levan, prior to (blue) or after (green) the incubation with *S*. Tm for 24 h. Data in E and F represent min-max-scaled, normalized peak area (or concentration [in g/L] for formate) averaged over 5 biological replicates. White crossed-out cells indicate that a metabolite was below the limit of detection in the respective sample.

To test whether *B. theta* PUL inhibition impacts *S.* Tm cross-feeding, we performed an adapted spent media assay (Figure 5B; Methods). Overnight cultures of *B. theta* wild-type, deletion, or complementation strains were pelleted, resuspended in M13 supplemented with the respective polysaccharide, and—informed by their previously determined growth kinetics (Suppl. Figure S8A-D)—incubated for either 4 h (levan, hyaluronic acid, polygalacturonic acid) or 8 h (dextran) at 37°C. The resulting supernatants were sterile-filtered, inoculated with *S.* Tm, and bacterial growth was monitored under aerobic conditions for 24 h. Strikingly, the growth of *S.* Tm in the supernatants of PUL22– and PUL48-inactivated mutants was strongly decreased compared to supernatants from both wild-type and complemented *B. theta* in levan or dextran, respectively (Figure 5C, D). In comparison, more subtle differences in *S.* Tm growth were observed in the supernatants of PUL57 and PUL75 mutants (Suppl. Figure S9B, C).

To pinpoint metabolites cross-fed via PUL22 and PUL48, we collected the supernatants of the respective *B*. *theta* strains before and after the incubation with *S*. Tm and analyzed them by targeted metabolomics (raw data in Suppl. Table 5). In dextran, the majority of *B*. *theta*-derived metabolites depended on intact PUL48 activity (Figure 5E, Suppl. Figure S10). Exceptions were mannitol/sorbitol, glutamine, and alanine, which accumulated independently of PUL48, as well as serine, isoleucine, and leucine, whose levels even increased when PUL48 was inactivated. Blocking PUL22-mediated levan utilization (Figure 5F, Suppl. Figure S11) led to reduced levels of several carbohydrates (including glucose, fructose, GalNAc, inulobiose, ketose, and melezitose), of most amino acids (except for glutamine and serine), and of SCFAs and their intermediates. The few remaining monosaccharides in the PUL22 mutant-derived supernatant were completely used up by *S*. Tm after 24 h, highlighting the massive impact of PUL22 for cross-feeding under this condition. Of note, succinate, glutamate, and aspartate—identified above as important substrates for *S.* Tm during sugar scarcity (Figure 4E, F)—were efficiently reduced by blocking either PUL48 or PUL22. Conversely, and reflecting *S*. Tm growth kinetics (Suppl. Figure S9B, C), only subtle metabolite differences were observed in analogous experiments with PUL57 and –75 mutants (Suppl. Figure S9D, E).

In summary, of four tested PULs, the inhibition of PUL22– and PUL48-specific polysaccharide degradation prevented the accumulation of usable substrates for *S.* Tm growth. We therefore focus on these two PULs from here on.

### Tweaking individual components of PUL22 to fine-tune B. theta–S. Tm cross-feeding

Deleting transcriptional PUL activators, as done above, silences the cognate PUL machinery. In contrast, selectively disrupting PUL-associated membrane proteins involved in substrate binding (SusD-like) and import (SusC-like) is expected to retain extracellular processing of polysaccharides by outer membrane-bound CAZymes, while preventing the import of the resulting oligosaccharides. In such a case, target polysaccharide utilization by *B. theta* would be selectively inhibited, but *S*. Tm cross-feeding would still occur or even be enhanced. To test this hypothesis, we focused on PUL22 and its substrate, levan, and generated a cognate *susD* knockout strain (Δ*susD^BT1762^*). As previously described (Sonnenburg *et al*., 2010), this mutant exhibited a growth defect in M13 supplemented with levan, suggesting that this single *susD* inactivation is sufficient to abrogate *B. theta* levan utilization (Suppl. Figure S8E). We note, however, that *susD^BT1762^ trans*-complementation failed to recover wild-type growth (again similar to previous observations (Sonnenburg *et al*., 2010)), presumably because complemented *BT1762* did not reach wild-type expression levels (Suppl. Figure S8J).

Incubating the Δ*susD^BT1762^ B. theta* mutant in levan yielded substantial amounts of free fructose and di-/trisaccharides, suggesting the release of the building blocks of levan by extracellular glycoside hydrolase (GH) activity (Figure 5F). Accordingly, *S*. Tm growth in Δ*susD^BT1762^*-derived supernatants was comparable to that from wild-type *B. theta* cultures and drastically increased compared to supernatants from the PUL22 null mutant (Δ*HTCS^BT1754^*) (Figure 5D). Metabolomics indicated the monosaccharides released by Δ*susD^BT1762^ B. theta* to be consumed by *S*. Tm, whereas the levels of di– and trisaccharides increased upon *S*. Tm growth (Figure 5F). A plausible explanation for the latter may be that *S*. Tm deploys enzymes to process fructan-derived oligosaccharides, resulting in the accumulation of smaller sugar chains in the supernatant. Since sugar molecules larger than trisaccharides evade detection by GC/LC-MS, this hypothesis could not be supported by our metabolomics data.

The surface-exposed GH BT1760 functions as an endo-levanase (Sonnenburg *et al*., 2010) and may thus contribute to cross-feeding from *B. theta* to *S.* Tm. We constructed strains overexpressing *BT1760* in either the wild-type or Δ*susD^BT1762^* background (Suppl. Figure S8K) and performed an adapted spent media assay in levan. In both cases, we observed enhanced pathogen growth relative to *B. theta* wild-type supernatants (Figure 5D). These results suggest an important role of this *B. theta* endo-levanase in liberating *S*. Tm-accessible sugars, both during unperturbed PUL22-driven levan degradation and when the import of levan-derived oligosaccharides into *B. theta* is blocked. This validates our hypothesis that *S*. Tm cross-feeding can be fostered by selectively interfering with the cognate oligosaccharide importers in *B. theta*. Overall, the targeted manipulation of individual components of this levan-catabolic PUL system affected *S.* Tm growth in a predictable manner, showcasing the adjustability of *B. theta*’s glycan degradation machinery for commensal-pathogen cross-feeding.

### Disrupting PUL48-mediated cross-feeding blunts Salmonella outgrowth in vivo

To investigate whether the PUL-dependent inhibition of cross-feeding *in vitro* would also counteract *S*. Tm growth under *in vivo* conditions, we utilized a gnotobiotic mouse model. We chose to focus on PUL48-dependent dextran utilization in the mouse model, as dextran, unlike many other polysaccharides, is readily soluble in water and does not increase viscosity, thereby not interfering with uptake by mice. Germ-free mice were pre-colonized with *B. theta* wild-type or a mutant strain devoid of the SusR-like activator of dextran-catabolic PUL48 (Δ*susR^BT3091^*) (Figure 6A). Following a pre-colonization period of seven days, mice were switched from chow to a semisynthetic diet (SSD), with one subgroup each receiving additional dextran in their drinking water. One day after the diet change, all mice were challenged with *S.* Tm ΔΔ*spi-1/2* (an avirulent *S.* Tm strain unable to invade host cells and cause an inflammatory response (Hapfelmeier *et al*., 2005; Stecher *et al*., 2007; Schulte *et al*., 2011)). From day –1 to day 3 post-infection (dpi), fecal samples were collected daily and colonization densities of *B. theta* and *S.* Tm were determined by selective plating. Longitudinal quantification of *B. theta* CFUs revealed an increase following the SSD switch, yet independent of dextran supplementation, in all treatment groups (Figure 6B, C). Furthermore, neither the dietary switch to SSD nor the additional dextran supplementation resulted in significant differences in colonization levels between *B. theta* wild-type and the Δ*susR^BT3091^* strain (Figure 6B, C; Suppl. Figure S12A). This implies that its metabolic flexibility renders *B. theta* relatively resistant to the inhibition of specific PULs, at least within this simplified host-associated model.

**Figure 6:**
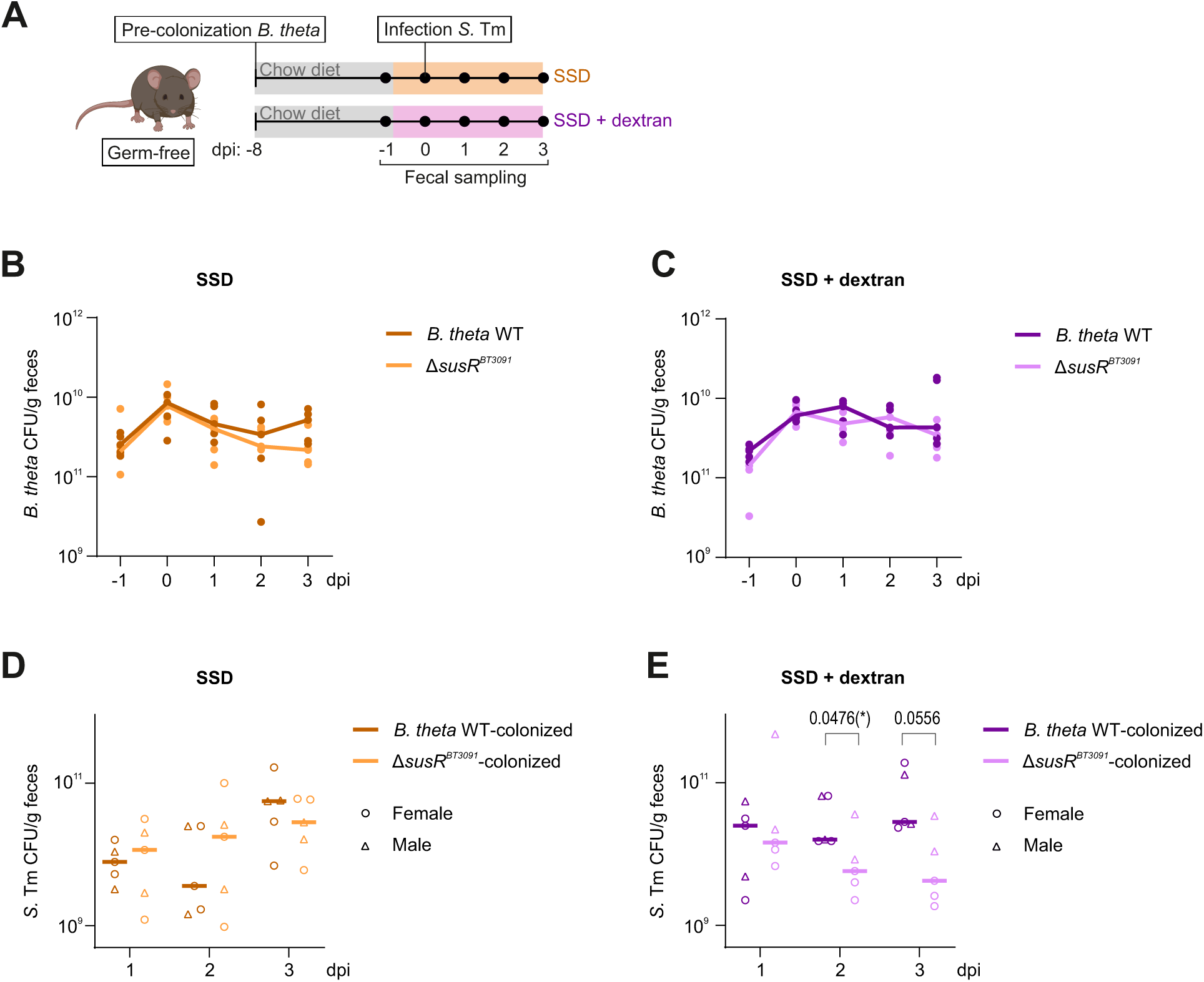
Disrupting PUL-dependent cross-feeding reduces *Salmonella* levels in the mouse gut. A: Experimental outline of cross-feeding experiment in gnotobiotic mice. B-C: CFU counts of *B. theta* wild-type and Δ*susR^BT3091^* in fecal pellets of mice fed a semi-synthetic diet (SSD) without (B) or with (C) dextran supplemented in the drinking water. D-E: CFU counts of *S*. Tm in fecal pellets of mice that had been pre-colonized with either wild-type or Δ*susR^BT3091^ B. theta* and fed the SSD diet, either without (D) or with (E) dextran in the drinking water. Data are differentially plotted for female (circles) and male (triangles) mice. A non-parametric Mann-Whitney test was performed to test for statistical significance in CFU differences between *B. theta* wild-type– and Δ*susR^BT3091^*-colonized mice at individual time points. The bold lines plotted in panels B-E refer to the median of 5 mice per group, each plotted as an individual dot. dpi: days post-infection.

Following *Salmonella* infection, the avirulent *S*. Tm strain expanded drastically within one day, with a 10^4^-to 10^5^-fold increase in CFUs compared to the input (10^5^ CFU/mouse). *S*. Tm densities between mice pre-colonized with *B. theta* wild-type or Δ*susR^BT3091^* did not differ on the three consecutive days post-infection in mice fed the SSD diet (Figure 6D). Importantly however, *S*. Tm CFUs in Δ*susR^BT3091^*-pre-colonized mice were significantly reduced on dpi 2 and 3 when the mice were additionally supplemented with dextran (Figure 6E). This effect was more pronounced in female than in male mice (Suppl. Figure S12B, C).

Collectively, these results suggest that cross-feeding from *B. theta* to *S.* Tm occurs in the murine intestine and that it can be prevented in a host environment by inhibiting specific polysaccharide degradation systems in *B. theta*.

## DISCUSSION

The degradation of plant– and host-derived polysaccharides by gut commensals is a central process that governs microbial interactions, shapes community structures, and profoundly contributes to host health. Consequently, targeting metabolic processes in the gut microbiota has been proposed as a possibility to interfere with enteric infection (Passalacqua, Charbonneau and O’Riordan, 2016; Jones, de Brito and Byndloss, 2025). Focusing on a model pair of gut commensal and pathogen, we set out to demonstrate causalities in carbon source cross-feeding.

Our work identified extensive nutrient handover from *B. theta* to *S.* Tm that encompasses carbohydrates, amino acids, and SCFAs, thereby complementing previous findings (Ng *et al*., 2013; Faber *et al*., 2017; Spiga *et al*., 2017; Shelton *et al*., 2022). Of these *B. theta*-released metabolites, mono– and disaccharides were the preferred substrates of *Salmonella*. However, a *S*. Tm mutant defective in glucose import was able to exploit alternative carbon sources, particularly succinate and the amino acids glutamate and aspartate (Fig. 4). This is in line with previous studies that demonstrated that *S*. Tm can utilize microbiota-derived succinate (Spiga *et al*., 2017) and that both, glutamate and aspartate, serve as critical intermediates feeding into *S*. Tm’s TCA cycle when glucose is scarce (Zampieri *et al*., 2019). In contrast, *B*. *theta* PUL inhibition not only reduced the levels of available simple sugars, but also efficiently depleted these alternative *S*. Tm carbon sources from the environment (Fig. 5E, F). Consequently, and contrasting the marginal effects of *S*. Tm glucose uptake inactivation, *B. theta* PUL inhibition entailed strong suppression of *Salmonella* growth (Fig. 5C, D). These findings support our initial hypothesis that the restriction of *Salmonella*-accessible nutrients, rather than interfering with metabolic processes in the pathogen itself, could represent an attractive anti-infective strategy. Furthermore, we demonstrated this suppressive effect for a specific PUL system and diet combination in a gnotobiotic mouse model. At least in the absence of metabolic competition with companion microbiota species, *B. theta*’s *in vivo* fitness was not negatively impacted by the inhibition of PUL48, even when the corresponding polysaccharide substrate (dextran) was supplemented to the host diet. While encouraging, more research is warranted before generalizing these findings.

High levels of monosaccharides in the colon are typically considered detrimental due to their association with irritable bowel syndrome, type 2 diabetes, colon cancer, and other diseases (Arnone *et al*., 2022). However, under certain conditions it may be desirable to transiently increase simple sugar levels in the colon, e.g. as part of a prebiotic regimen to stimulate their fermentation into host-beneficial metabolites such as butyrate (Bedu-Ferrari *et al*., 2022). In this context it is worth reiterating that our study provided proof-of-principle that *Bacteroides* PULs can be predictably manipulated in both directions (Fig. 5): inhibiting their activity to decrease product accumulation, but also—by selectively blocking SusCD-mediated sugar import—increase the levels of available polysaccharide degradation products in the surrounding.

Apart from these potentially clinically exploitable findings, this study provides insight into the regulatory principles of glycan degradation in *B*. *theta*. Our comprehensive gene expression data of this key gut symbiont in the presence of diverse, *in vivo*-relevant polysaccharides builds on previous efforts to decode *B*. *theta*’s transcriptomic landscape (Martens, Chiang and Gordon, 2008; Ryan *et al*., 2020, 2024; Rüttiger *et al*., 2025) and were integrated in our Theta-Base browser for easy access by the public (https://bacteroides.helmholtz-hzi.de/jbrowse_new). In combination with the extensive polysaccharide-dependent metabolome analyses, these data represent a rich resource to help improve our functional understanding of catabolic processes in this gut commensal. As one example, PUL enrichment analysis revealed a few, simple PULs to be induced by glucose rather than by any complex glycan. It is intriguing that these PULs lack a sensory/regulatory component, which suggests that they are controlled by external, *trans*-acting factors, such as transcriptional regulators of non-cognate PULs (Lynch and Sonnenburg, 2012), global metabolic transcription factors such as Cur (Schwalm, Townsend and Groisman, 2016; Townsend *et al*., 2020), or post-transcriptionally, through small regulatory RNAs or RNA-binding proteins (Ryan *et al*., 2020; Rüttiger *et al*., 2025). Indeed, expression of one such PUL (PUL18; *BT1551-BT1553*) was previously found to be reduced in a Cur-deficient *B. theta* mutant compared to wild-type during growth on glucose (Townsend *et al*., 2020), suggesting PUL18 to be governed by Cur in a sugar-responsive manner. Further exploring such atypical PULs could reveal novel regulatory networks mediating carbon prioritization in *Bacteroides*.

Taken together, this study dissects the molecular underpinnings of *Bacteroides* PUL-mediated metabolite cross-feeding to *Salmonella*. The obtained findings improve our understanding of the basic principles of polysaccharide catabolism and nutrient sharing in the gut. It remains to be seen to what extent this knowledge may help inform strategies to manipulate the indigenous microbiota to the host’s benefit.

### Limitations of the study

This proof-of-principle study deliberately focused on the molecular basis of cross-feeding between only two select bacterial species. Clearly, translating the presented findings into the complex gut environment, characterized by a diverse community of commensal species and a fluctuating supply of different carbon sources, represents a formidable challenge. For example, we did not address here to what extent the discovered principles would also account for commensal *Enterobacteriaceae* of the gut microbiota that compete with invading enterobacterial pathogens for nutrients (Osbelt *et al*., 2024; Wende *et al*., 2025). Additionally, our study focused on unilateral cross-feeding, while accumulating evidence highlights the existence of bidirectional metabolic relationships in the intestinal microbiota (Huus *et al*., 2021). Relevant in the present context, *B*. *theta* hijacks iron-laden siderophores from *Enterobacteriaceae* to acquire this essential cofactor in the intestine (Zhu *et al*., 2020). In the future, repeating some of the presented experiments in the additional presence of commensal *Escherichia coli* species—alone or within a simplified microbial community (syncom)—would help decipher these interactions and represent a practical intermediate step towards a conventional microbiota. The latter also comprises numerous other *Bacteroides* species that might compensate for metabolites otherwise derived from *B*. *theta* PUL activity. In this regard, antisense oligonucleotide (ASO) technology may come in handy, for it is highly programmable and can be multiplexed (Vogel, 2020). While classically used as antimicrobials, ASOs could be devised to transiently shut down specific catabolic processes, not only in *B*. *theta*, but in multiple *Bacteroides* spp. in parallel, to rationally manipulate the bulk metabolic profile of the microbiome.

## SUPPLEMENTARY FIGURE LEGENDS

**Suppl. Figure 1:**
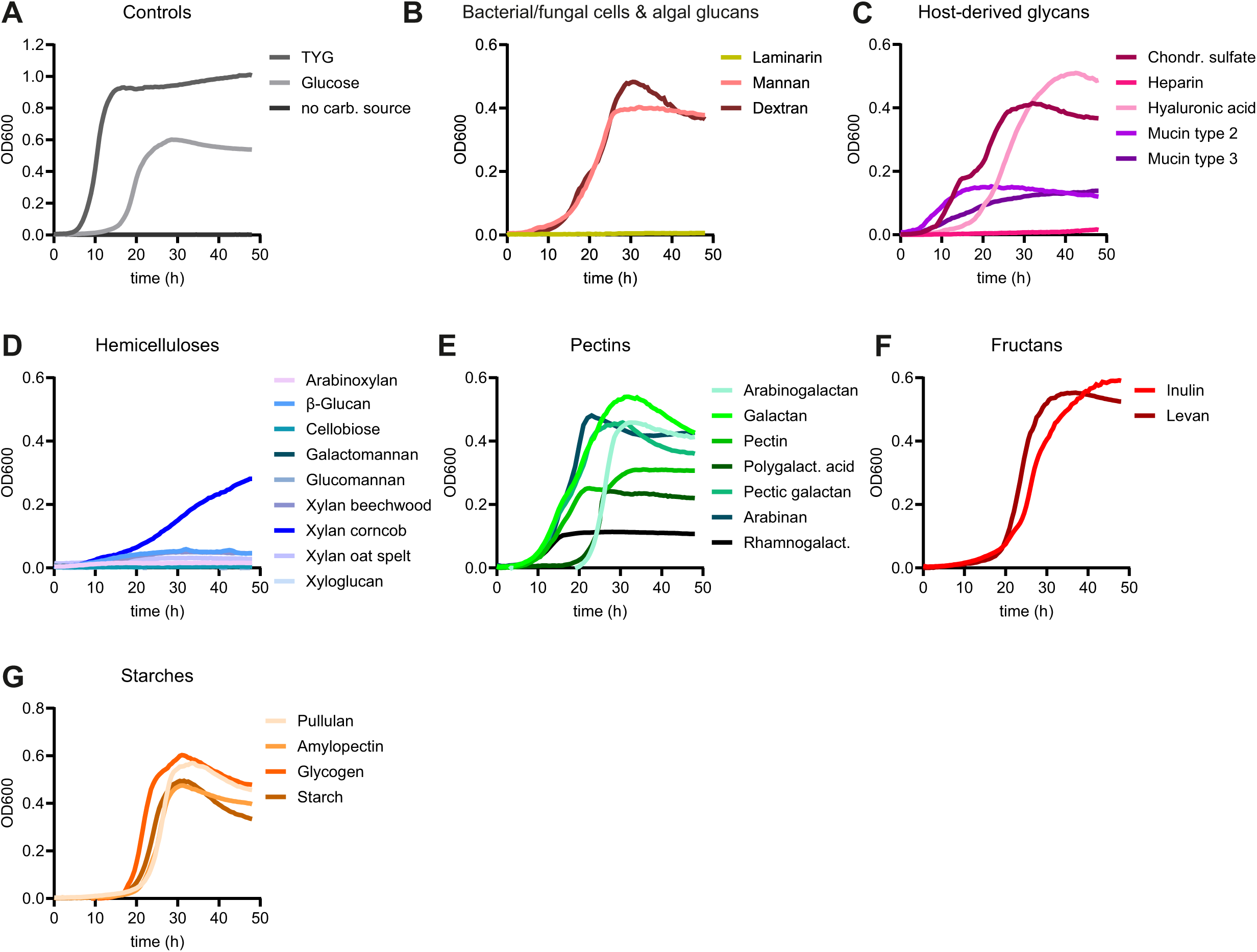
Growth curves of *B. theta* in VB medium supplemented with different glycans. A-G: *B. theta* grown in two different control media (VB medium with glucose, TYG) or in the absence of a carbon source (A) and in VB medium supplemented with polysaccharides belonging either to the group of bacterial/fungal cell walls and algal glucans (B), host-derived glycans (C), hemicelluloses (D), pectins (E), fructans (F), or starches (G). Graphs show the average growth from ≥2 biological replicates, each including three technical replicates.

**Suppl. Figure 2:**
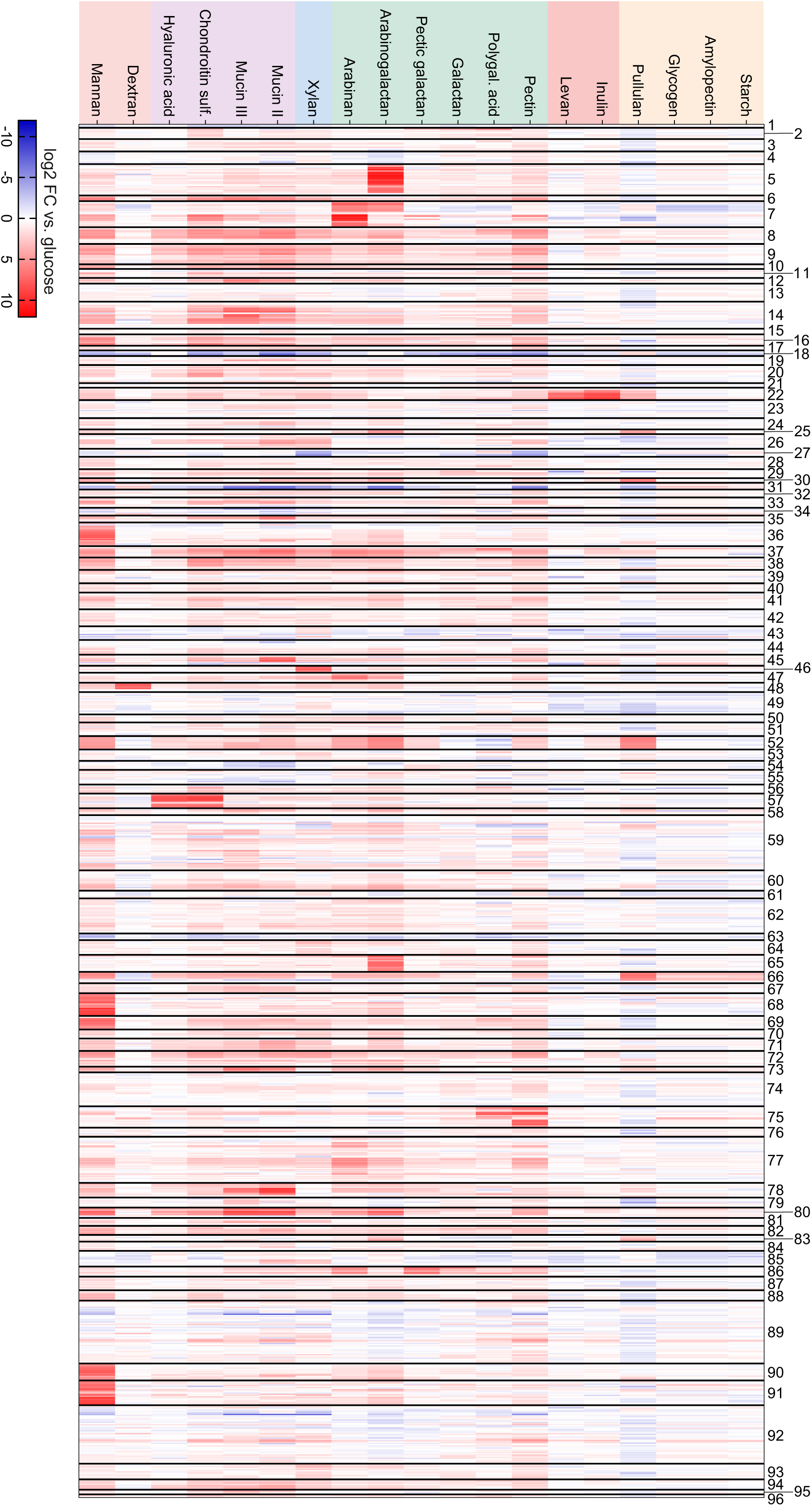
Expression of *B. theta* PUL-associated genes during growth on different polysaccharides. Heatmap depicts the mean log2 fold-changes of each PUL gene from *B. theta* grown in VB medium supplemented with the respective polysaccharide relative to glucose. Mean values were calculated from 2 biological replicates. Numbers to the right refer to the literature-derived PUL nomenclature.

**Suppl. Figure 3:**
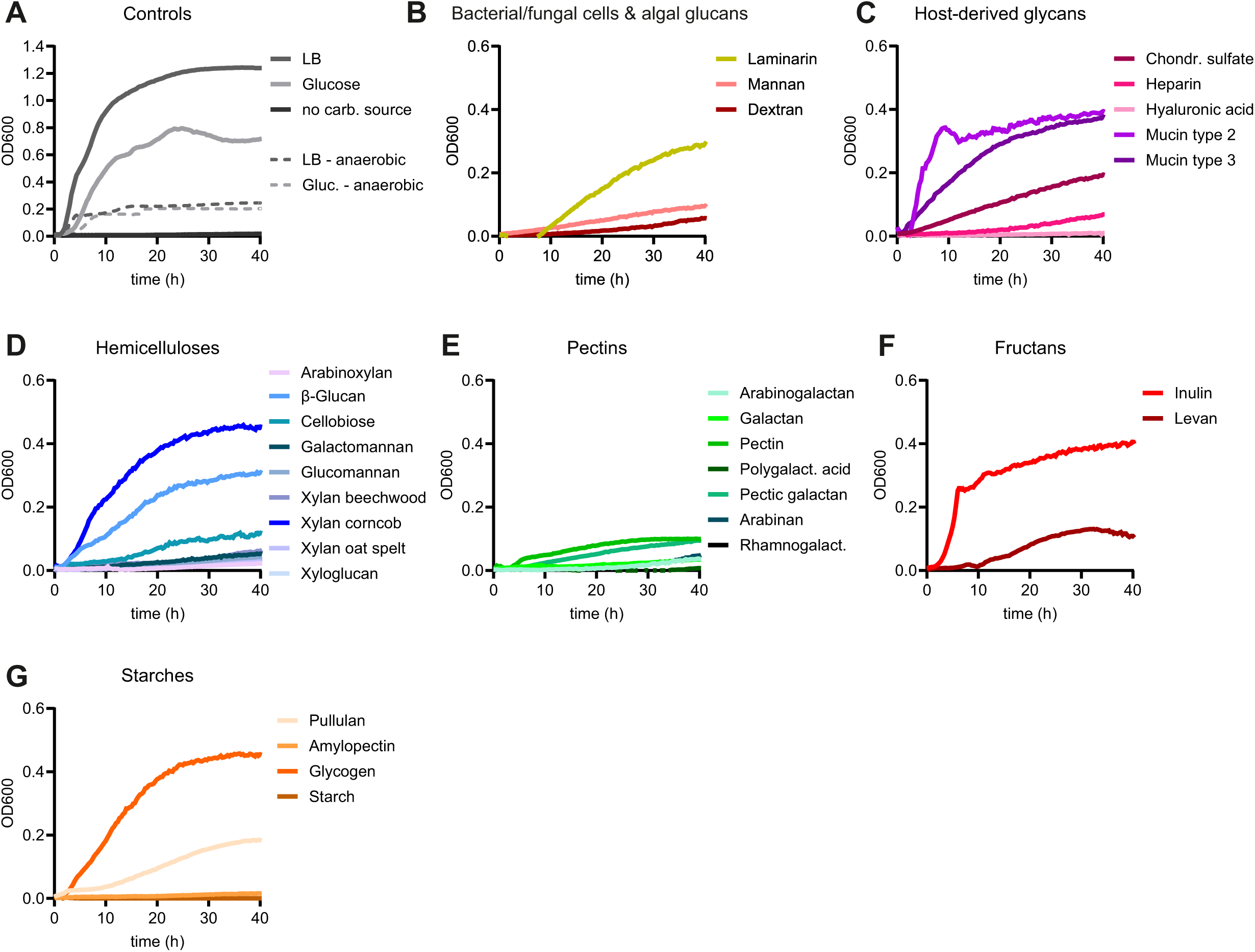
Growth curves of *S.* Tm in M9 medium supplemented with different glycans. A-G: Aerobic (solid lines) and anaerobic (dashed lines) growth of *S*. Tm in control media (M9 medium with glucose, LB) or in the absence of a carbon source (A) and aerobic growth of *S*. Tm in M9 medium supplemented with polysaccharides belonging to the groups of bacterial/fungal cell walls and algal glucans (B), host-derived glycans (C), hemicelluloses (D), pectins (E), fructans (F), and starches (G). Graphs show the average growth from ≥ 2 biological replicates, each including three technical replicates.

**Suppl. Figure 4:**
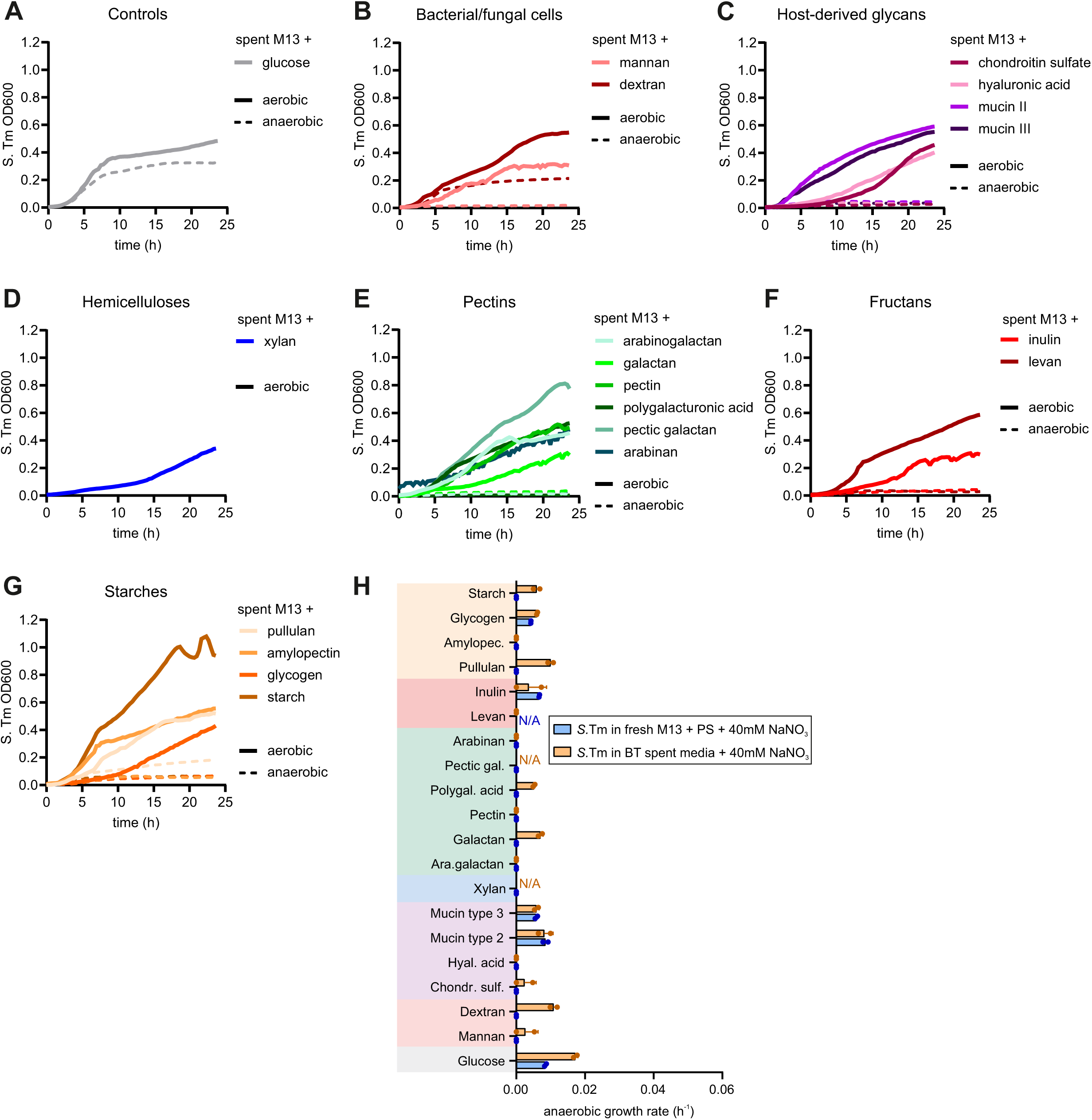
Growth kinetics of *S.* Tm in spent medium from *B. theta*. A-G: Aerobic (solid lines) and anaerobic (dashed lines) growth of *S*. Tm in *B. theta*-processed M13 medium supplemented with glucose (A) or with complex carbon sources belonging to the groups of bacterial/fungal cell walls and algal glucans (B), host-derived glycans (C), hemicelluloses (D), pectins (E), fructans (F), and starches (G). Graphs depict the mean growth calculated from five biological replicates under aerobic conditions or two biological replicates under anaerobic conditions. H: Anaerobic growth rates of *S.* Tm in fresh M13 supplemented with the indicated polysaccharides (blue bars) or in sterilized supernatants from *B. theta* that had been grown in the same media (orange bars), each supplemented with 40 mM NaNO_3_. Each dot represents one biological replicate, comprised of each three technical replicates.

**Suppl. Figure 5:**
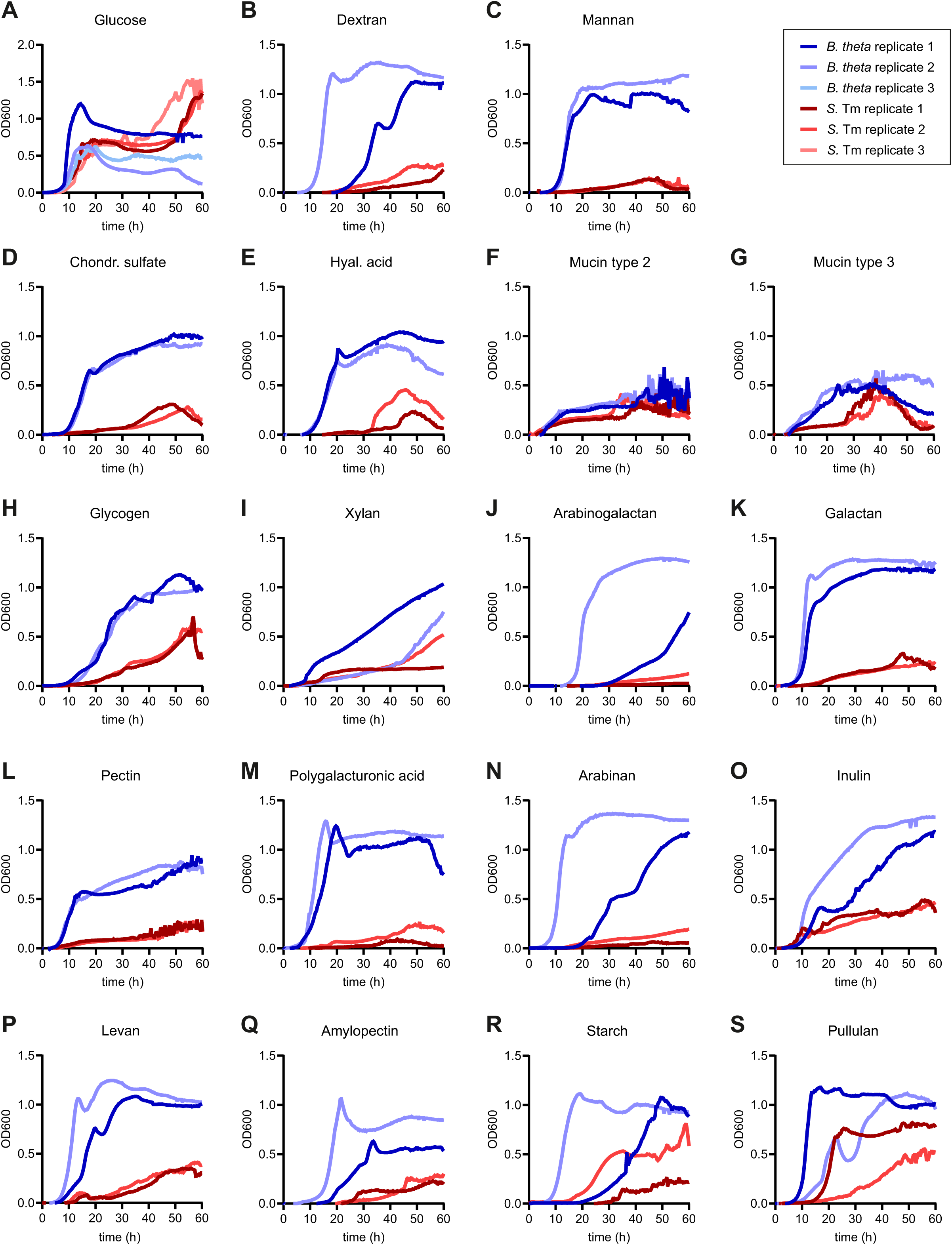
Growth curves of *B. theta* and *S.* Tm in a co-culture assay under anaerobic conditions. A-S: Growth of *B. theta* (blue) and *S.* Tm (red) monitored in parallel in co-culture plates filled with M13 supplemented with glucose (A) or with 18 different plant-or host-derived polysaccharides (B-S). The data reflect the average from 2-3 biological replicates per carbon source.

**Suppl. Figure 6:**
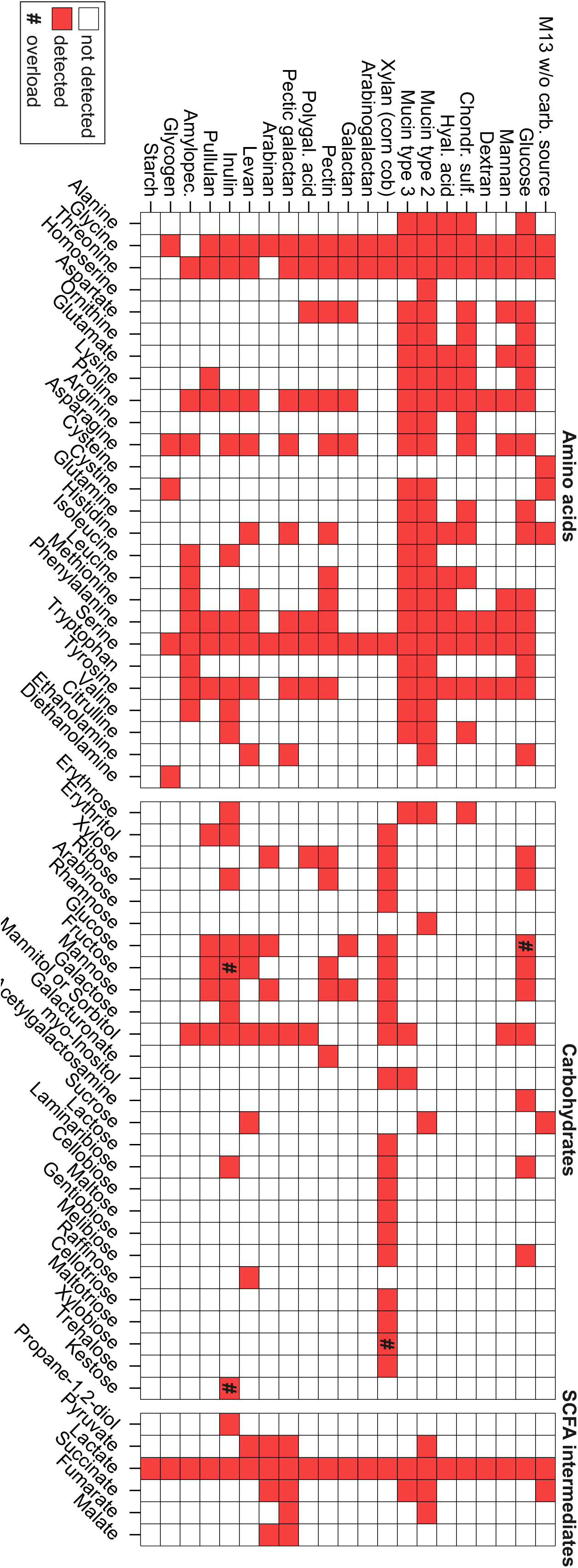
Quantification of free amino acids, carbohydrates, and SCFA intermediates detected in blank and polysaccharide-containing M13 medium. Heatmap visualizes the presence (red) or absence (white) of the indicated metabolites, grouped by chemical class, across the 19 polysaccharide-containing media and glucose and carbon source-free M13 media as reference. The hash symbol indicates overload for the respective metabolite and sample.

**Suppl. Figure 7:**
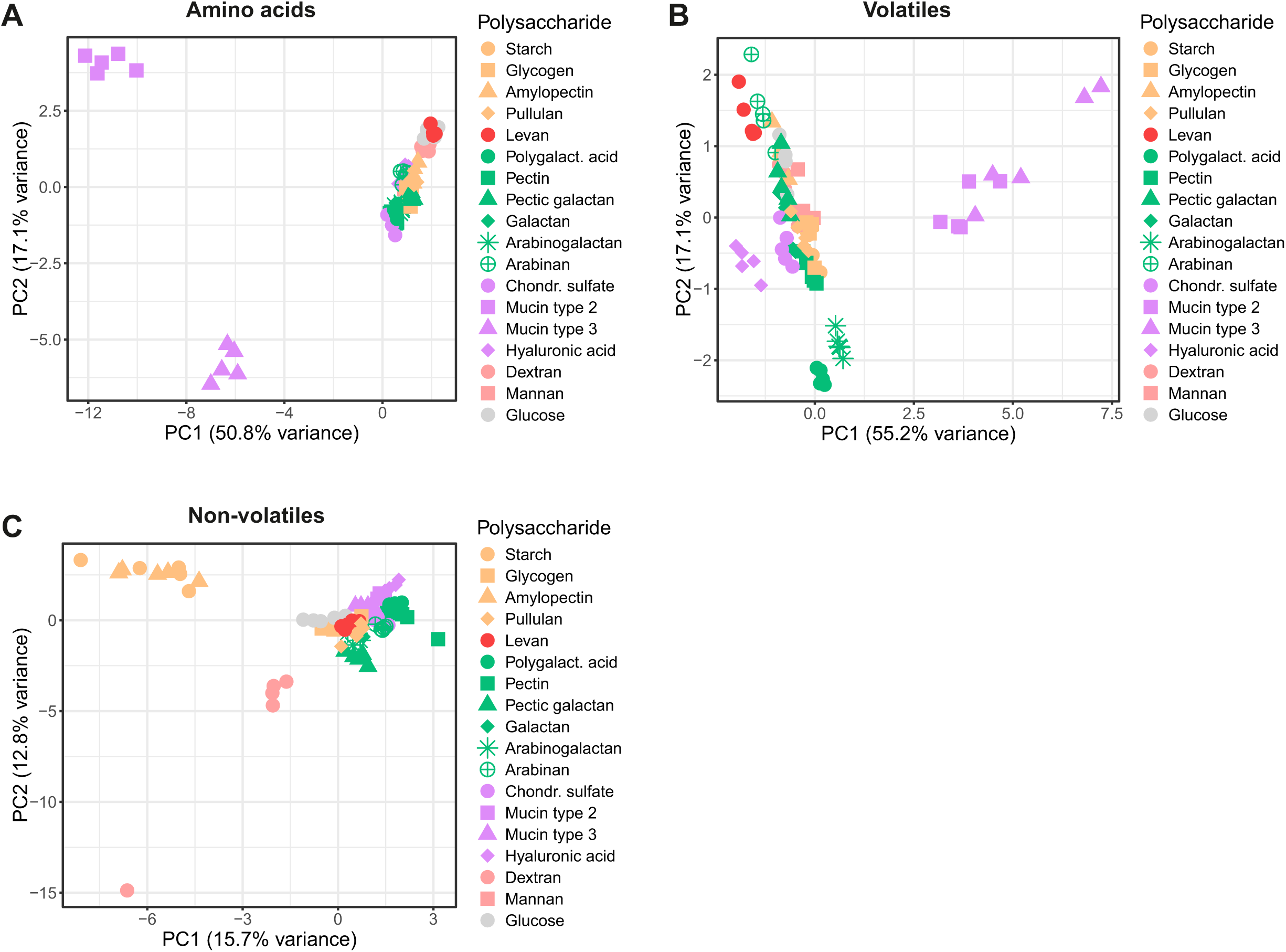
Metabolome profiles of *B. theta* grown in M13 supplemented with various polysaccharides. A-C: PCoA plots of total amino acids (A), volatiles (B), and non-volatiles (C) detected in the supernatant of *B. theta* cultures grown in M13 supplemented with different polysaccharides and glucose as control. Each data point represents one biological replicate and colors reflect different polysaccharide classes.

**Suppl. Figure 8:**
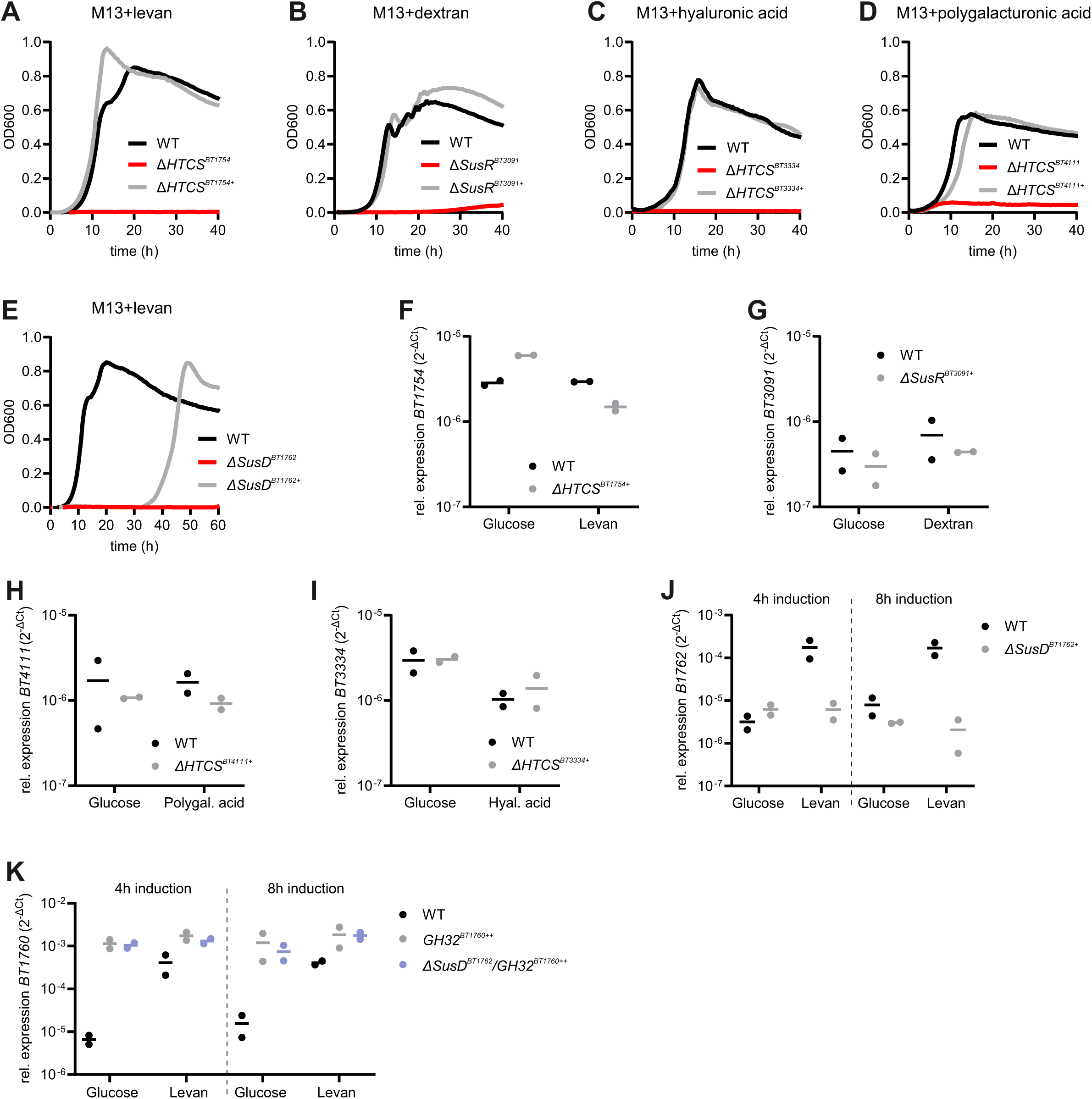
Growth curves of *B. theta* wild-type, PUL regulator mutants, and *trans*-complementation strains. A: Growth of *B. theta* wild-type, Δ*HTCS^BT1754^*, and Δ*HTC^SBT1754+^* in M13 supplemented with levan. B: Growth of *B. theta* wild-type, Δ*susR^BT3091^*, and Δ*susR^BT3091+^* in M13 supplemented with dextran. C: Growth of *B. theta* wild-type, Δ*HTCS^BT3334^*, and Δ*HTCS^BT3334+^* in M13 supplemented with hyaluronic acid. D: Growth of *B. theta* wild-type, Δ*HTCS^BT4111^*, and Δ*HTCS^BT4111+^* in M13 supplemented with polygalacturonic acid. E: Growth of *B. theta* wild-type, Δ*susD^BT1762^*, and Δ*susD^BT1762+^* in M13 supplemented with levan. The curves plotted in panels A-E represent mean values calculated from three technical replicates. F-I: Relative expression of *BT1754* (F), *BT3091* (G), *BT4111* (H), and *BT3334* (I) in *B. theta* wild-type or *trans*-complemented strains in M13 medium with either glucose or levan, dextran, polygalacturonic acid or hyaluronic acid, as determined by qRT-PCR. J: Relative expression of *BT1762* in *B. theta* wild-type or Δ*susD^BT1762+^* after 4 or 8 h of incubation in M13 with either glucose or levan, as determined by qRT-PCR. K: Relative expression of *BT1760* in *B. theta* wild-type and *GH32^BT1760^*-overexpression strains (the latter, either in the wild-type or Δ*susD^BT1762^* background) after 4 or 8 h of incubation in M13 with glucose or levan, as determined by qRT-PCR. Panels F-K: each dot represents one biological replicate, calculated from each three technical replicates.

**Suppl. Figure 9:**
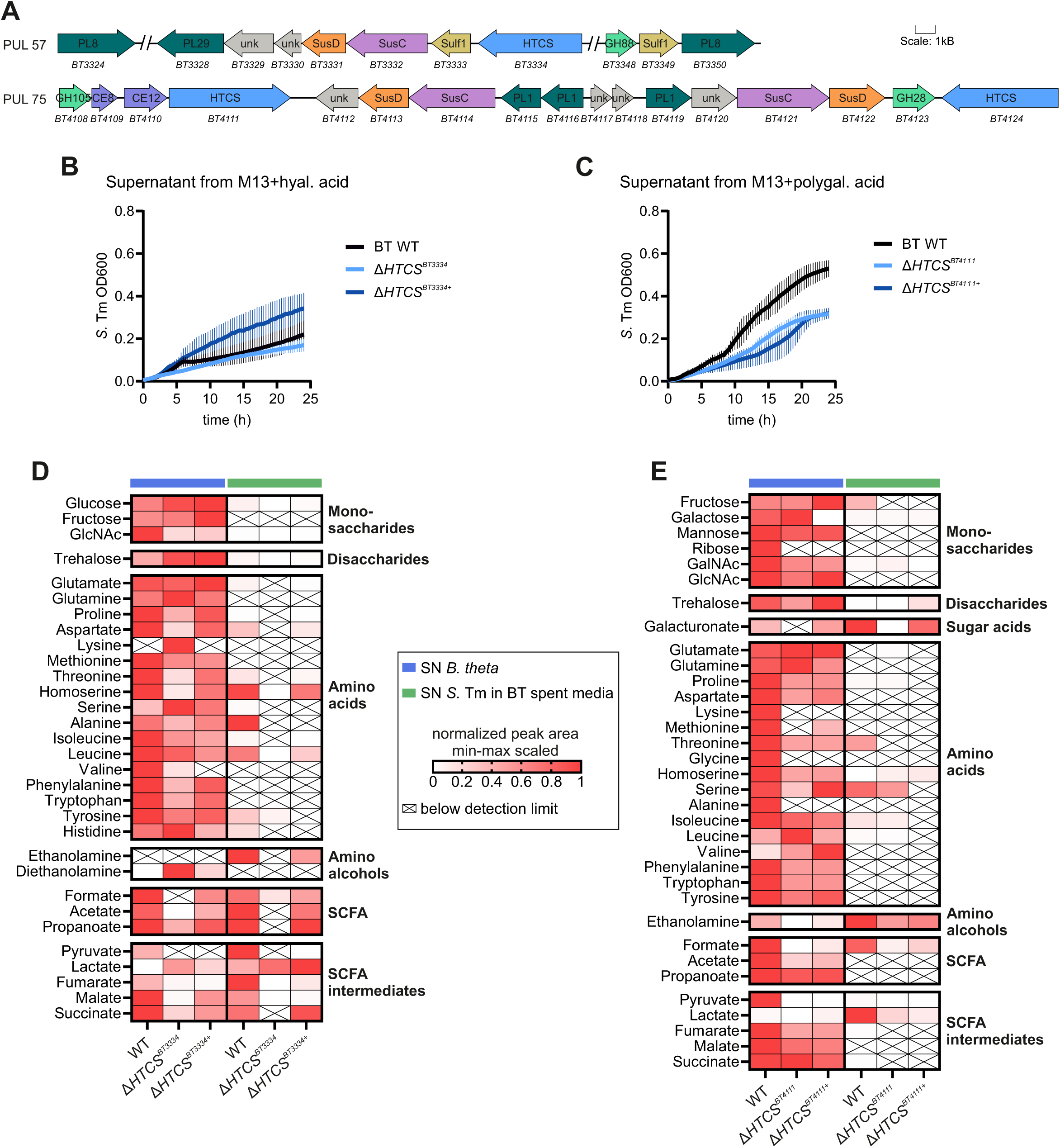
Mild effects of the genetic inactivation of *B. theta* PUL57 and PUL75 on *S*. Tm cross-feeding. A: Schematic of *B. theta* PULs 57 and 75. Locus tags and protein functions derived from cazy.org. HTCS: hybrid two-component system, GH: glycoside hydrolase, PL: polysaccharide lyase, Sulf: sulfatase, CE: carbohydrate esterase, unk: unknown (hypothetical protein). B: Growth of *S.* Tm in supernatants from *B. theta* wild-type, Δ*HTCS^BT3334^*, or complementation strains incubated for 4 h in M13 with hyaluronic acid. C: Growth of *S.* Tm in supernatants from *B. theta* wild-type, Δ*HTCS^BT4111^*, or complementation strains incubated for 4 h in M13 with polygalacturonic acid. D: Metabolites measured in the supernatants of *B. theta* wild-type, Δ*HTCS^BT3334^*, or complementation strains grown on hyaluronic acid (blue rows) and after growth of *S.* Tm in the *B. theta*-derived supernatants (green rows). E: Metabolites measured in the supernatants of *B. theta* wild-type, Δ*HTCS^BT4111^*, or complementation strains grown in polygalacturonic acid (blue rows) or after growth of *S.* Tm in the *B. theta*-derived supernatants (green rows). Data in D and E represent the min-max-scaled normalized peak area (or, in case of formate, the concentration in g/L) averaged over 5 biological replicates. White crossed-out cells indicate that a metabolite was below the detection limit in the respective sample.

**Suppl. Figure 10:**
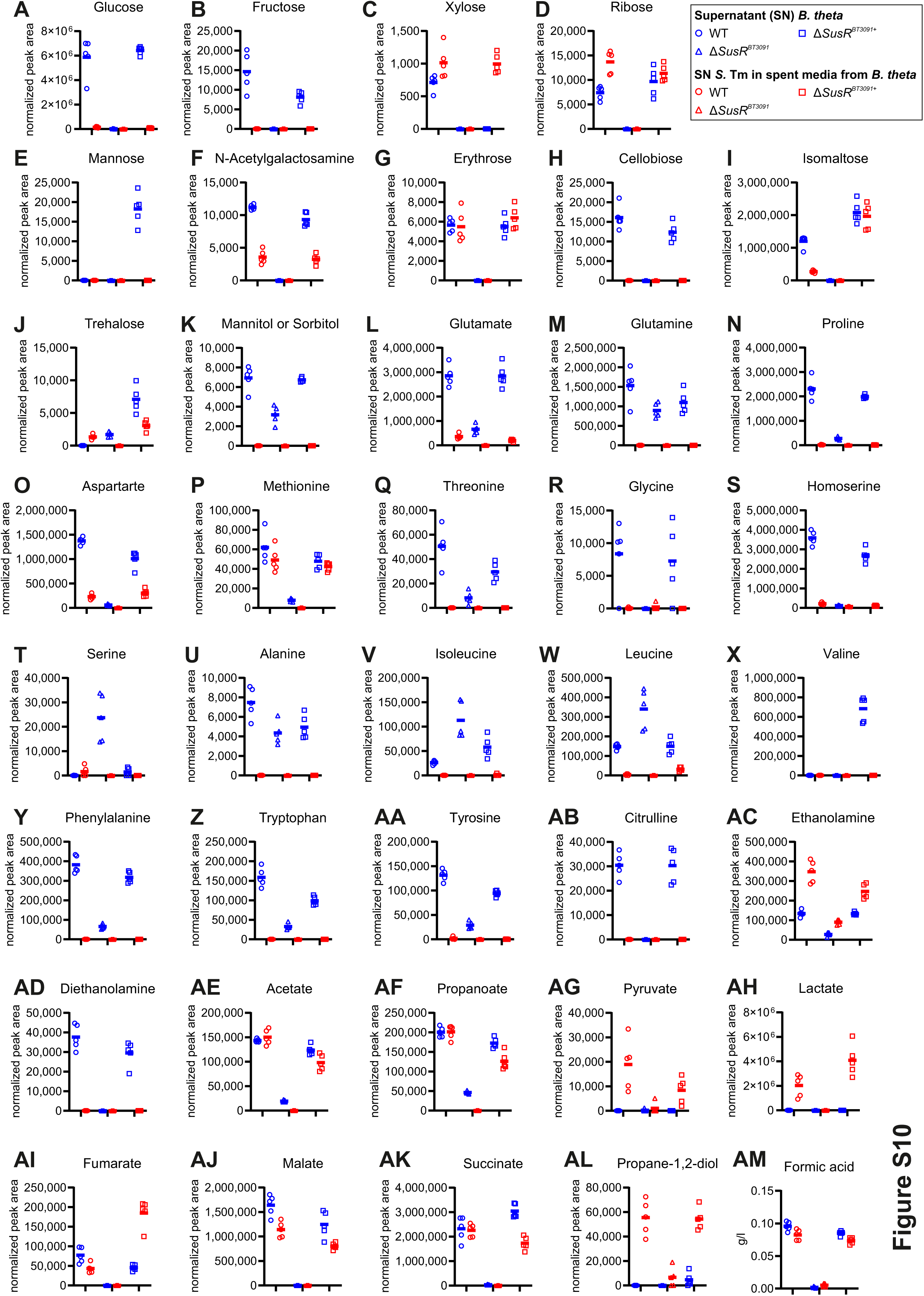
Metabolite quantification in supernatants from different *B. theta* PUL48 strains and *S.* Tm in M13 with dextran. Normalized peak areas (A-AL) and concentration (g/L) (AM) of metabolites measured in the supernatants from *B. theta* wild-type, Δ*susR^BT3091^*, and Δ*susR^BT3091+^* in M13 containing dextran, before and after 24 h of growth of *S.* Tm.

**Suppl. Figure 11:**
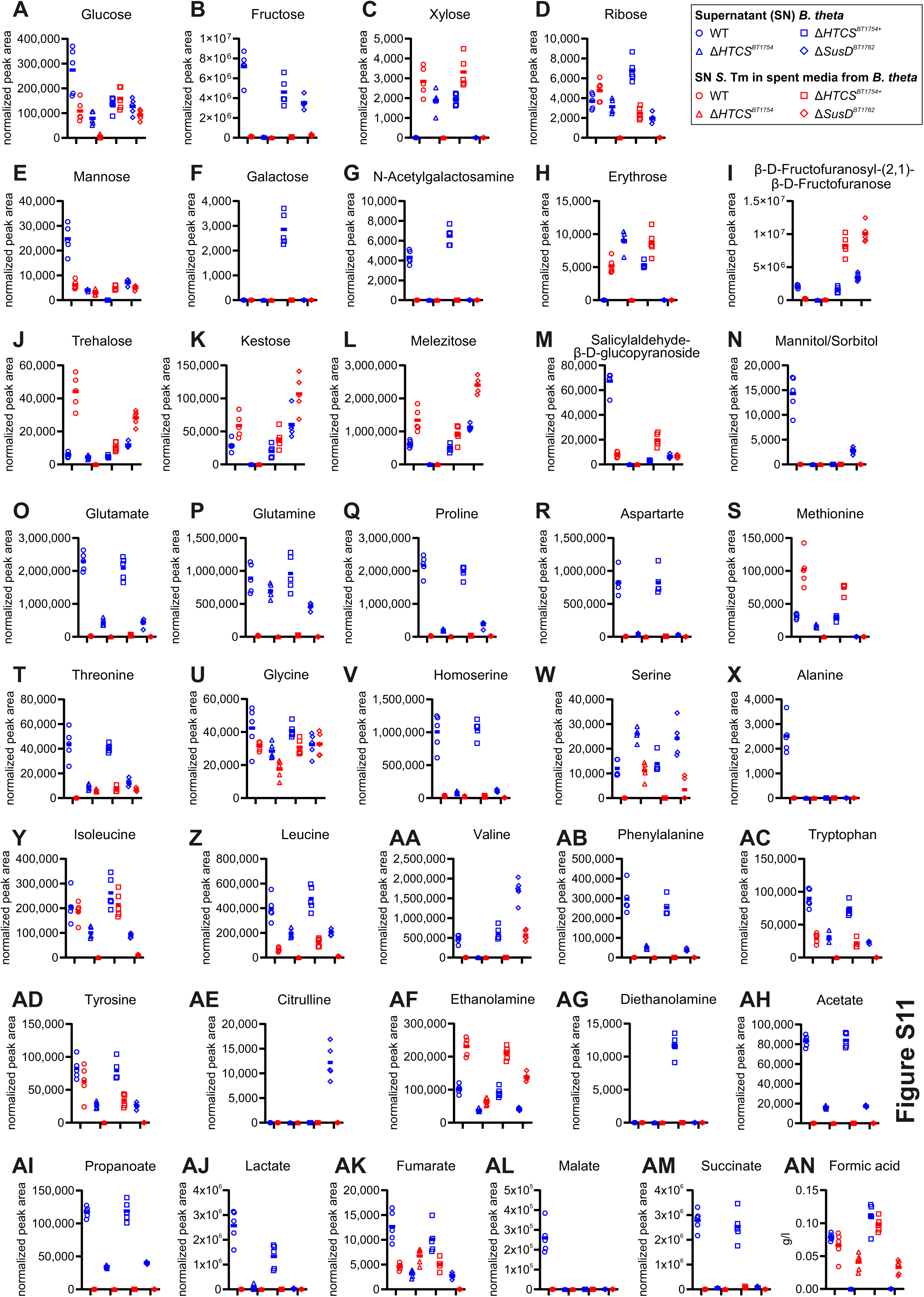
Metabolite quantification in supernatants from different *B. theta* PUL22 strains and *S.* Tm in M13 with levan. Normalized peak areas (A-AM) and concentration (g/L) (AN) of metabolites measured in the supernatants from *B. theta* wild-type, Δ*HTCS^BT1754^*, Δ*HTCS^BT1754+^*, and Δ*susD^BT1762^* in M13 containing levan, before and after 24 h of growth of *S.* Tm.

**Suppl. Figure 12:**
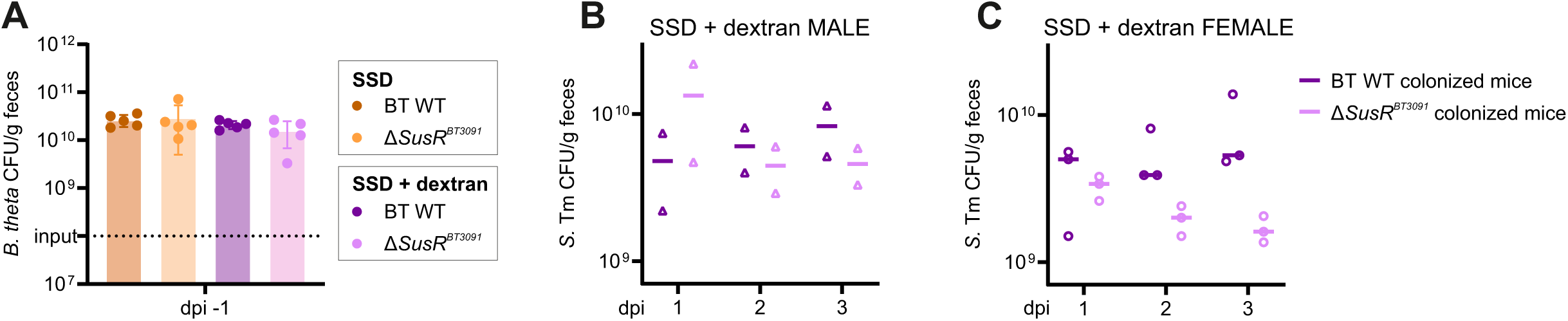
*B. theta* and *S.* Tm CFU counts from the cross-feeding experiment in gnotobiotic mice. A: *B. theta* CFUs in the feces of mice seven days after initial colonization via oral gavage and prior to *S.* Tm infection (dpi –1). B-C: *S.* Tm CFUs in the feces of male (B) and female mice (C) pre-colonized with either *B. theta* wild-type or the PUL48-associated Δ*susR^BT3091^* mutant and fed with a semi-synthetic diet (SSD) supplemented with dextran in the drinking water. Panels A-C: each data point refers to an individual mouse.

## SUPPLEMENTARY TABLES

**Suppl. Table 1: Overview of complex carbohydrates used in this study.**

**Suppl. Table 2: Sampling ODs for RNA–seq experiment and spent media assay.**

**Suppl. Table 3: Metabolite data of the spent media assay.**

**Suppl. Table 4: Metabolite data of spent media assay with *S*. Tm wild-type and *S*. Tm ΔΔΔ*ptsG/manXYZ/glk* in *B. theta* spent M13 + glucose or dextran**

**Suppl. Table 5: Metabolite data of the adapted spent media assay with *B. theta* wild-type, deletion mutants, and complementation strains in dextran, levan, hyaluronic acid, and polygalacturonic acid.**

## METHODS

### Bacterial strains and growth conditions

The following bacterial strains were used in this study: *Bacteroides thetaiotaomicron* VPI-5482, *Salmonella enterica* serovar Typhimurium SL1344, both from DSMZ-German Collection of Microorganisms and Cell Cultures GmbH; *S*. Tm ΔΔ*spi-1/2*, provided by Marc Erhardt (Humboldt University Berlin, Germany); *S.* Tm ATCC 14028 and *S.* Tm ΔΔΔ*ptsG*/*manXYZ*/*glk*, provided by Thilo Fuchs (Friedrich Loeffler Institute, Jena, Germany). *B. theta* strains were routinely grown in TYG media (20 g/L tryptone, 10 g/L yeast extract, 0.5% glucose, 5mg/l hemin, 1 g/L L-cysteine, 0.0008% CaCl_2_, 19.2 mg/L MgSO_4_ x 7H_2_O, 40 mg/L KH_2_PO_4_, 40 mg/L K_2_HPO_4_, 80 mg/L NaCl, 0.2% NaHCO_3_) or BHI-S plates (52 g/L BHI agar, 1 g/L cysteine, 5 mg/L hemin, 0.2% NaHCO_3_) at 37°C in an anaerobic chamber from Coy Laboratories with a gas mixture composition of 10% H_2_, 10% CO_2_, 80% N_2_. *S.* Tm was routinely grown in LB media (5 g/L yeast extract, 5 g/L NaCl, 10 g/L tryptone) or LB plates (5 g/L yeast extract, 5 g/L NaCl, 10 g/L tryptone, 18 g/L agar) in 37°C under aerobic conditions.

The following minimal media compositions were used in this study: VB medium (1 g/l L-cysteine, 5 mg/L hemin, 20 mg/l L-methionine, 4.17 mg/l FeSO_4_, 0.2% NaHCO_3_, 0.9 g/L KH_2_PO_4_, 0.02 g/l MgCl_2_ x 6H_2_O, 0.026 g/L CaCl_2_ x 2H_2_O, 0.001 g/L CoCl_2_ x 6H_2_O, 0.01 g/L MnCl_2_ x 4H_2_O,

0.5 g/L NH_4_Cl, 0.25 g/L Na_2_SO_4_) (Varel and Bryant, 1974); M9 medium (0.004% Histidine, 0.5 µg/mL Thiamine, 0.1 mM CaCl_2_, 2 mM MgSO_4_, 12,8 g Na_2_HPO_4_ x 7H_2_O, 3 g KH_2_PO_4_, 0.5 g NaCl, 1 g NH_4_Cl) (Miller 1972); M13 medium (1.2 mg/L Hematin, 31 mg/L L-Histidine, 0.5 g/L L-Cysteine, 0.384 mg/L FeSO_4_ x 7H_2_O, 9.52mg/l MgCl_2_, 5.55mg/l CaCl_2_ (anhydrous), 0.88g/l NaCl, 1.12 g/L (NH_4_)2SO_4_, Vitamin B12 5 µg/L, Vitamin K3 1mg/l, KH_2_PO_4_ 13.6g/l) (Tramontano *et al*., 2018). Minimal media were either supplemented with 5 g/L glucose or 5 g/L of the respective polysaccharides, which were resuspended in water and autoclaved prior to use.

### Gene editing in *B. theta*

Gene deletion in *B. theta* was performed as previously described using the pSIE1 plasmid system (Bencivenga-Barry *et al*., 2020). In brief, 750 bp flanking regions up– and downstream of the gene of interest were assembled into the pSIE vector by using the NEBuilder HiFi DNA Assembly kit. The resulting plasmid was electroporated into *E. coli* S17-1 λpir and subsequently conjugated into *B. theta.* Selection of the conjugants were performed by plating on BHI-S+gentamycin+erythromycin plates. Colonies were inoculated into TYG medium and the resulting culture was plated in serial dilutions on BHI-S plates supplemented with anhydrotetracycline. Colonies were screened by PCR and further confirmed by Sanger sequencing. Gene complementation was performed by cloning the gene of interest together with its native promoter sequence into the plasmid pWW3452 (Whitaker, Shepherd and Sonnenburg, 2017) using the NEBuilder HiFi DNA Assembly kit. Overexpression of a gene was achieved analogously by replacing the native promoter with a strong phage promoter. For both variants, the plasmid was transformed into *E. coli* S17-1 λpir, conjugated into the *B. theta* deletion mutant and were selected on BHI-S+gentamycin+erythromycin plates.

### Growth curves

A single colony was inoculated into 5 mL of minimal media supplemented with 5 g/L glucose. Following a 24 h incubation, 1 mL of the culture was pelleted (3,000 g, 3 min) and the pellet was resuspended in PBS. 200 µL of fresh minimal media supplemented with different polysaccharides (5 g/L) was loaded into wells of a 96-well plate and was inoculated in a 1:100 dilution with the previously prepared PBS suspension. Absorbance at 600 nm was measured at 20-30 min intervals using a Biotek plate reader.

### RNA extraction

Bacterial cultures for RNA isolation were harvested in early stationary phase by mixing a volume corresponding to an optical density (OD) equivalent of 4 with 20% stop mix (95% ethanol, 5% phenol, pH > 7.0) and snap-freezing in liquid nitrogen. Isolation of RNA was performed by hot-phenol extraction. In brief, the bacterial suspension was thawed on ice, pelleted and cells were lysed by adding 600 µL lysozyme (0.5 mg/mL) and 60 µL of 10% SDS followed by a 2 min incubation at 64°C and further addition of 66 µL 3 M NaOAc. After that, 750 µL phenol were added, the mixture was incubated for 6 min at 64°C and was subsequently centrifuged. The aqueous phase was transferred to a PLG heavy tube, 750 µL chloroform was added and followed by additional centrifugation. RNA was precipitated by mixing the aqueous phase with twice the volume of a 30:1 mixture ethanol:3 M NaOAc (pH 6.5) and overnight incubation at –20°C. Afterwards, samples were centrifuged, the resulting pellet was washed with 75% ethanol and was eventually resuspended in RNase-free water. Contaminating DNA was removed by treatment with DNase I (5 U per 40 µg RNA) and 0.5 µL of RNase Inhibitor for 45 min at 37°C. RNA was purified by another phenol-chloroform extraction followed by ethanol precipitation and resuspension of the pellet in RNase-free water.

### RNA sequencing-based transcriptome analysis

RNA sequencing, read processing, and mapping was performed as previously described (Ryan *et al*., 2024). In brief, RNA samples were depleted of ribosomal RNA, treated with DNase, and libraries were prepared. Following cDNA synthesis, sequencing was performed on the NextSeq 500 platform by the SysMed Core Unit at the University of Würzburg. Generated reads were quality checked, adapters were cut and reads were mapped to the *B. theta* VPI-5482 reference genome (NC_004663.1).

Differential gene expression analysis was performed as previously described (Ryan *et al*., 2024) with comparing gene expression during growth in polysaccharides with gene expressing during growth in glucose as control. PCoA to visualize global gene expression profiles was performed using read counts obtained from the trimmed DESeq2 result tables. To identify significantly enriched PUL gene sets in the respective carbohydrate conditions, gene set enrichment analysis was performed using the *fgsea* (v1.28.0) package in R studio. PUL gene sets were retrieved from the CAZy database (cazy.org). Normalized enrichment scores were calculated and PULs with a FDR (adjusted p-value) < 0.05 were considered significantly enriched.

### Spent media assay

A single colony of *B. theta* was inoculated into 5 ml of M13 supplemented with 5 g/L glucose. Following a 24 h incubation, 1 mL of the culture was pelleted (3,000 g, 3 min), the pellet was resuspended in PBS and OD-adjusted. M13 supplemented with 5 g/L of the respective polysaccharide was inoculated with bacterial suspension (1:100 dilution) and was incubated for 16-20 h depending on the growth rate of *B. theta* in the respective polysaccharide. Resulting cultures were centrifuged (4,700 g, 10 min, RT) and resulting supernatant was sterile filtered using filters with 0.2 µm pore size. Supernatants were dispensed into wells of a 96-well plate and inoculated with *S.* Tm. The latter was previously grown from a singly colony in M13 plus glucose for 24 h, pelleted, resuspended in PBS and OD adjusted. Bacterial growth of inoculated supernatants were assessed under aerobic and anaerobic conditions using a Biotek reader and measuring absorbance at 600 nm every 20 min for a total of 24 h.

### Co-culture assay

Single colonies of *B. theta* and *S.* Tm were grown individually in M13 plus glucose under anaerobic conditions for 24 h. The resulting pre-culture was pelleted (3,000 g, 3 min), the pellet was resuspended in PBS and OD was adjusted to 1.0. Each side of a co-culture duet (Cerillo) was filled with 700 µL of M13 supplemented with the respective polysaccharide and was either inoculated with the OD-adjusted *B. theta* or *S.* Tm pre-culture (1:100 dilution). Bacterial growth in each duet was measured every 20 min under anaerobic conditions for 60 h.

### Adapted spent media assay

Single colonies of *B. theta* wild-type, deletion or complementation strain were inoculated into M13 plus glucose and were grown for 24 h under anaerobic conditions. The resulting cultures were subcultured into M13 plus glucose and were grown for another 16 h. The OD was adjusted and 5 mL of the resulting culture was pelleted by centrifugation (10 min, 4,700 rpm). The resulting pellet was resuspended in M13 with the respective polysaccharide and was incubated anaerobically for 4 h (regulator mutants and complementation strains in levan, hyaluronic acid, and polygalacturonic acid) or 8 h (regulator mutants and complementation strains in dextran as well as glycoside hydrolase overexpression strain in levan). After this incubation, the cultures were centrifuged again (10 min, 4,700 rpm, 4°C) and 1.5 mL of the resulting supernatant was aliquoted and snap frozen in liquid nitrogen for subsequent metabolome analysis. The remaining supernatant was sterile filtered using 0.2 µm filters. The supernatant was used to perform (i) growth curves with *S.* Tm (as described above) and (ii) metabolome analysis after *S.* Tm growth. For that, 2ml of the supernatant was inoculated 1:100 with *S.* Tm, which was previously grown from single colonies in M13 plus glucose for 24 h, pelleted, resuspended in PBS and OD-adjusted. Following aerobic incubation for 24h, the culture was pelleted (centrifugation 10 min, 4,700 rpm, 4°C) and 1.5 mL of the supernatant were snap frozen for metabolome analysis.

### Metabolite detection and analysis

*GC/MS measurement of non-volatile metabolites:* Samples were diluted 1:10 with water. 50 µL of the dilution were mixed with 30 µL of ^13^C-ribitol solution (4% of ribitol stock solution [0.2 mg/mL] in methanol) as internal standard. Samples were dried in in a vacuum concentrator. Metabolite analysis was performed on an Agilent GC-MSD system (7890B coupled to a 5977 GC) equipped with a high-efficiency source and a Gerstel RTC system, as published previously (Will *et al*., 2019). Briefly, following a two-step, automated derivatization with a methoxyamine hydrochloride solution and N-methyl-N-(trimethylsilyl)-trifluoracetamide, 1 µL of the sample was injected into a multimode inlet in pulsed splitless mode and pulsed split mode with a split ratio of 10:1. Separation was conducted on an Agilent VF-5ms column with a helium flow (1.2 mL/min). Data were analyzed using the MetaboliteDetector Software (Hiller *et al*., 2009, version 2.2N). Chromatographic peak area (based on selected quantification ions) of each measured compound was normalized to the peak area of ^13^C-ribitol as internal standard. Normalized peak areas were used for further analysis.

*GC/MS measurement of short chain fatty acids:* Samples were diluted 1:10 with water. 400 µL of the dilution were used for an ether extraction and further mixed with 600 µL of H_2_O/internal standard (o-cresol, 8 µL/L), 60 µL of H_2_SO_4_ (MS grade), and 200 µL of methyl tert-butyl ether (MTBE). Mixtures were vortexed for 5 min and centrifuged for 5 min at maximum speed. The ether phase was analyzed by GC-MS as previously described (Neumann-Schaal *et al*., 2015), using an Agilent GC-MSD system (7890B coupled to a 5977 GC) equipped with a high-efficiency source (Agilent Technologies, CA, USA). Data were analyzed using the MetaboliteDetector Software (Hiller *et al*., 2009). Chromatographic peak area (based on selected quantification ions) of each measured compound was normalized to the peak area of o-cresol as internal standard. Normalized peak areas were used for further analysis.

*LC/MS measurement of amino acids:* Samples were diluted 1:10 with water and 50 µL were mixed with 5 µL of internal standard solution (sarcosine 0.044 mg/mL, norvaline 0.058 mg/mL). Samples were analyzed using an Intrada amino acid column (3 x 100 mm, 3 µm, Imtakt Corp., Kyoto, Japan) according to the technical note TI839E of Imtakt with some modifications: solvent A (acetonitrile:water 60:40 with 0.5% formic acid) and solvent B (acetonitrile:water 20:80, 80 mM ammonium formate) were used starting with 85% solvent A/15% solvent B for 4 min and a gradient to 100% solvent B for 3 min followed by 100% solvent B for 5 min at a flow rate of 0.6 mL/min. Amino acids were detected using an Agilent HPLC-MS system of an 1290 Infinity II series HPLC coupled to an 6545 QTOF equipped with a dual AJS ESI source in positive mode. Due to the detection limit, glycine was analyzed as part of the non-volatile metabolites (see above). Chromatographic peak area (based on the protonated adduct) of each measured compound was normalized to the peak area of norvaline and sarcosine as internal standards. For determination of the retention times, an Amino Acid Standard (5061-3330, Agilent Technologies, CA, USA) supplemented with asparagine, glutamine, ornithine, and citrulline in a concentration range of 5-200 µM was used. Data were analyzed using the Agilent MassHunter Qualitative Navigator software (version B.08.00) and normalized peak areas were used for further analysis.

*Formate quantification:* Formate was analyzed using the Megazyme Formic Acid Kit (Megazyme, Bray, Ireland) according to the microplate protocol provided by the manufacturer. Absolute concentrations (g/L) were determined based on a standard calibration control.

Metabolite concentrations based on normalized peak areas of blank samples (M13 supplemented with the respective polysaccharide) were subtracted from all bacterial supernatant samples. For PCoA analysis (Figure 3A, B; Suppl. Figure 7), below threshold values were imputed using the minimal detected value for the respective metabolite multiplied by factor 0.95. Standardization across different metabolites was achieved by calculation z-scores, which standardizes the original data based on the mean and standard deviation of each metabolite data. For heatmap visualization (Figure 3C-E; 5E, F; Suppl. Figure S9D, E), the arithmetic mean of the blank-corrected values from the five biological replicates was calculated for each condition. These mean values were then min-max scaled separately for each metabolite. For metabolite logFCs (Figure 4E, F), log-transformed fold changes were calculated by comparing normalized peak area values (non-blanked) from ΔΔΔ*ptsG*/*manXYZ*/*glk* to wild-type *S.* Tm (ATCC 14028) across replicate pairs.

### Quantitative real-time PCR

qRT-PCR was performed from two biological replicates in technical triplicates. For each sample, 5 μL master mix (Takyon No Rox SYBR Master Mix dTTP Blue) 0.1 μL forward primer (10 μM), 0.1 μL reverse primer (10 μM), 0.1 μL reverse transcriptase and 3.7 μL RNase-free water were mixed with 1 μL of DNase-treated RNA (10 µg/µL). qRT-PCR was performed in a CFX96 Touch Real-Time PCR Detection System. Gene expression was analyzed using the 2−ΔCt method.

### Mouse work

Germ-free C57BL/6NTac mice were bred in isolators in the GF facility at the Helmholtz Centre for Infection Research, Braunschweig, Germany. Mice were kept under a strict 12 h light cycle and were co-housed in groups of 2-5 mice per cage. Sterilized food and water were available to all animals ad libitum.

For colonization of mice with *B. theta*, strains were grown overnight under anaerobic conditions in 5 mL TYG. The next morning, the culture was pelleted by centrifugation at 3,000 g for 3 min, supernatant was removed and the pellet was resupended in PBS. OD was measured, adjusted, and mice were gavaged with 5 x 10^8^ CFU/mL. For infection with *S.* Tm, 25 mL of LB was inoculated with 1 mL of an *S.* Tm overnight culture and was incubated for 4 h at 37°C with constant shaking at 80 rpm. Following this, the culture was centrifuged for 15 min at 500 g, supernatant was removed and the pellet was resuspended in PBS. OD was adjusted and mice were gavaged with 5 x 10^5^ CFU/mL.

Colonization levels of *B. theta* and *S.* Tm were determined from fresh fecal samples of mice. Fecal samples were weighed and mixed with 0.5 mm zirconia/silica beads and 1 mL of sterile PBS. Samples were homogenized for 50 s at 2,400 rpm using a BioSpec Mini-Beadbeater-96. The resulting suspension was used to prepare dilution series. 25 µL of 10^0^–10^7^ dilutions were plated on BHI-S plates supplemented with 200 µg/mL gentamicin and were incubated anaerobically for 48 h to determine *B. theta* CFUs. In addition, 25 µL each were plated on LB plates supplemented with 100 µg/mL ampicillin and were incubated aerobically for 20 h to determine *S.* Tm CFUs.

### Statistical analyses

Statistical analyses were conducted in GraphPad Prism (v10.5.0) or R studio (v2023.12.1) using the packages ggplot2 (v3.5.1) stringr (v1.5.1), fgsea (v1.28.0), tidyverse (v2.0.0), scales (v1.3.0). The following statistical tests were used: Kolmogorov-Smirnov test combined with Benjamini-Hochberg FDR correction; paired, non-parametric Wilcoxon matched-pairs signed rank test; and non-parametric Mann-Whitney test. Description of the applied statistical test and P value thresholds of single experiments are given in the respective figure legends.

### Data availability

Raw sequencing data on GEO

Raw data of the metabolite analyses are deposited at the FAIRDOMHub

## ACKNOWLEDGEMENT

We are grateful to Marc Erhard, Thilo Fuchs, and Andreas Götz for *Salmonella* strains and thank the Core Unit Systems Medicine of the University of Würzburg and the Animal Facility of the HZI for assistance with RNA-seq and mouse experiments, respectively. We further thank Daniel Ryan and Gianluca Prezza for their support with the transcriptome analysis as well as Tom Guest for help with GEO data submission. This work was funded by the European Research Council (ERC Starting Grant #101040214 to A.J.W.) and by the German Research Foundation (DFG) through the SFB1583 DECIDE (#492620490; project A04 to A.J.W.). T.S. was supported by the DFG (EXC 2155; project number 390874280).

## REFERENCES

1. Aguirre, M. et al. (2016) ‘Diet drives quick changes in the metabolic activity and composition of human gut microbiota in a validated in vitro gut model’, Research in Microbiology, 167(2), pp. 114–125. Available at: 10.1016/J.RESMIC.2015.09.006.

2. Anderson, K.L. and Salyers, A.A. (1989a) ‘Biochemical evidence that starch breakdown by Bacteroides thetaiotaomicron involves outer membrane starch-binding sites and periplasmic starch-degrading enzymes’, Journal of Bacteriology, 171(6), pp. 3192–3198. Available at: 10.1128/jb.171.6.3192-3198.1989.

3. Anderson, K.L. and Salyers, A.A. (1989b) ‘Genetic evidence that outer membrane binding of starch is required for starch utilization by Bacteroides thetaiotaomicron’, Journal of Bacteriology, 171(6), pp. 3199–3204. Available at: 10.1128/jb.171.6.3199-3204.1989.

4. Arnone, D. et al. (2022) ‘Sugars and Gastrointestinal Health’, Clinical Gastroenterology and Hepatology, 20(9), pp. 1912–1924.e7. Available at: 10.1016/j.cgh.2021.12.011.

5. Arumugam, M. et al. (2011) ‘Enterotypes of the human gut microbiome’, Nature, 473(7346), pp. 174–180. Available at: 10.1038/nature09944.

6. Barrett, E.L. and Riggs, D.L. (1982) ‘Evidence for a Second Nitrate Reductase Activity That Is Distinct from the Respiratory Enzyme in Salmonella typhimurium’, Journal of Bacteriology, 150(2), p. 563. Available at: 10.1128/JB.150.2.563-571.1982.

7. Becker, D. et al. (2006) ‘Robust Salmonella metabolism limits possibilities for new antimicrobials’, Nature, 440(7082), pp. 303–307. Available at: 10.1038/nature04616.

8. Bedu-Ferrari, C. et al. (2022) ‘Prebiotics and the Human Gut Microbiota: From Breakdown Mechanisms to the Impact on Metabolic Health’, Nutrients, 14(10). Available at: 10.3390/NU14102096.

9. Bencivenga-Barry, N.A. et al. (2020) ‘Genetic Manipulation of Wild Human Gut Bacteroides’, Journal of Bacteriology. Edited by L.E. Comstock, 202(3), pp. 1–12. Available at: 10.1128/JB.00544-19.

10. Bjursell, M.K., Martens, E.C. and Gordon, J.I. (2006) ‘Functional Genomic and Metabolic Studies of the Adaptations of a Prominent Adult Human Gut Symbiont, Bacteroides thetaiotaomicron, to the Suckling Period’, Journal of Biological Chemistry, 281(47), pp. 36269–36279. Available at: 10.1074/JBC.M606509200.

11. Bowden, S.D. et al. (2014) ‘Nutritional and Metabolic Requirements for the Infection of HeLa Cells by Salmonella enterica Serovar Typhimurium’, PLOS ONE, 9(5), p. e96266. Available at: 10.1371/JOURNAL.PONE.0096266.

12. Culp, E.J. and Goodman, A.L. (2023) ‘Cross-feeding in the gut microbiome: Ecology and mechanisms’, Cell Host and Microbe, 31(4), pp. 485–499. Available at: 10.1016/j.chom.2023.03.016.

13. Cummings, J.H. et al. (1987) ‘Short chain fatty acids in human large intestine, portal, hepatic and venous blood’, Gut, 28(10), pp. 1221–1227. Available at: 10.1136/GUT.28.10.1221.

14. Cuskin, F. et al. (2015) ‘Human gut Bacteroidetes can utilize yeast mannan through a selfish mechanism’, Nature, 517(7533), pp. 165–169. Available at: 10.1038/nature13995.

15. Drula, E. et al. (2021) ‘The carbohydrate-active enzyme database: functions and literature’, Nucleic Acids Research, 50(D1), p. D571. Available at: 10.1093/NAR/GKAB1045.

16. Eisenreich, W. et al. (2010) ‘Carbon metabolism of intracellular bacterial pathogens and possible links to virulence’, Nature Reviews Microbiology, 8(6), pp. 401–412. Available at: 10.1038/NRMICRO2351.

17. Eng, S.K. et al. (2015) ‘Salmonella: A review on pathogenesis, epidemiology and antibiotic resistance’, Frontiers in Life Science, 8(3), pp. 284–293. Available at: 10.1080/21553769.2015.1051243.

18. Faber, F. et al. (2017) ‘Respiration of Microbiota-Derived 1,2-propanediol Drives Salmonella Expansion during Colitis’, PLoS Pathogens, 13(1), pp. 1–19. Available at: 10.1371/journal.ppat.1006129.

19. Ferreyra, J.A. et al. (2014) ‘Gut Microbiota-Produced Succinate Promotes C. difficile Infection after Antibiotic Treatment or Motility Disturbance’, Cell Host & Microbe, 16(6), pp. 770–777. Available at: 10.1016/J.CHOM.2014.11.003.

20. Foley, M.H., Cockburn, D.W. and Koropatkin, N.M. (2016) ‘The Sus operon: a model system for starch uptake by the human gut Bacteroidetes.’, Cellular and molecular life sciences: CMLS, 73(14), pp. 2603–17. Available at: 10.1007/s00018-016-2242-x.

21. Frolova, M.S. et al. (2022) ‘Genomic reconstruction of short-chain fatty acid production by the human gut microbiota’, Frontiers in Molecular Biosciences, 9, p. 862. Available at: 10.3389/FMOLB.2022.949563/BIBTEX.

22. Gennis, R.B., Stewart, V. and Neidhardt, F.C. (1996) ‘Escherichia coli and Salmonella: cellular and molecular biology’, Washington, DC, USA: American Society for Microbiology, pp. 217–261.

23. Götz, A. and Goebelt, W. (2010) ‘Glucose and glucose 6-phosphate as carbon sources in extra– and intracellular growth of enteroinvasive Escherichia coli and Salmonella enterica’, Microbiology, 156(4), pp. 1176–1187. Available at: 10.1099/mic.0.034744-0.

24. Han, J. et al. (2024) ‘Infection biology of Salmonella enterica’, EcoSal Plus, 12(1). Available at: 10.1128/ECOSALPLUS.ESP-0001-2023.

25. Hapfelmeier, S. et al. (2005) ‘The Salmonella Pathogenicity Island (SPI)-2 and SPI-1 Type III Secretion Systems Allow Salmonella Serovar typhimurium to Trigger Colitis via MyD88-Dependent and MyD88-Independent Mechanisms’, The Journal of Immunology, 174(3), pp. 1675–1685. Available at: 10.4049/jimmunol.174.3.1675.

26. Harrison, K.J., Crécy-Lagard, V. De and Zallot, R. (2017) ‘Gene Graphics: a genomic neighborhood data visualization web application’, Bioinformatics, 34(8), p. 1406. Available at: 10.1093/BIOINFORMATICS/BTX793.

27. Hiller, K. et al. (2009) ‘Metabolite detector: Comprehensive analysis tool for targeted and nontargeted GC/MS based metabolome analysis’, Analytical Chemistry, 81(9), pp. 3429–3439. Available at: 10.1021/AC802689C/ASSET/IMAGES/LARGE/AC-2008-02689C_0006.JPEG.

28. Huang, Y.L. et al. (2015) ‘Sialic acid catabolism drives intestinal inflammation and microbial dysbiosis in mice’, Nature Communications 2015 6:1, 6(1), pp. 1–11. Available at: 10.1038/ncomms9141.

29. Huus, K.E. et al. (2021) ‘Cross-feeding between intestinal pathobionts promotes their overgrowth during undernutrition’, Nature Communications, 12(1), pp. 1–14. Available at: 10.1038/s41467-021-27191-x.

30. Jones, K., de Brito, C.B. and Byndloss, M.X. (2025) ‘Metabolic tug-of-war: Microbial metabolism shapes colonization resistance against enteric pathogens’, Cell Chemical Biology, 32(1), pp. 46–60. Available at: 10.1016/j.chembiol.2024.12.005.

31. Kaoutari, A. El et al. (2013) ‘The abundance and variety of carbohydrate-active enzymes in the human gut microbiota’, Nature Reviews Microbiology 2013 11:7, 11(7), pp. 497–504. Available at: 10.1038/nrmicro3050.

32. Lapébie, P. et al. (2019) ‘Bacteroidetes use thousands of enzyme combinations to break down glycans’, Nature Communications 2019 10:1, 10(1), pp. 1–7. Available at: 10.1038/s41467-019-10068-5.

33. Liu, H. et al. (2021) ‘Functional genetics of human gut commensal Bacteroides thetaiotaomicron reveals metabolic requirements for growth across environments’, Cell Reports, 34(9), p. 108789. Available at: 10.1016/J.CELREP.2021.108789.

34. Louis, P. and Flint, H.J. (2017) ‘Formation of propionate and butyrate by the human colonic microbiota’, Environmental Microbiology, 19(1), pp. 29–41. Available at: 10.1111/1462-2920.13589.

35. Luis, A.S. et al. (2018) ‘Dietary pectic glycans are degraded by coordinated enzyme pathways in human colonic Bacteroides’, Nature Microbiology, 3(2), pp. 210–219. Available at: 10.1038/s41564-017-0079-1.

36. Lynch, J.B. and Sonnenburg, J.L. (2012) ‘Prioritization of a plant polysaccharide over a mucus carbohydrate is enforced by a Bacteroides hybrid two-component system’, Molecular Microbiology, 85(3), pp. 478–491. Available at: 10.1111/j.1365-2958.2012.08123.x.

37. Martens, E.C. et al. (2011) ‘Recognition and Degradation of Plant Cell Wall Polysaccharides by Two Human Gut Symbionts’, PLoS Biol, 9(12), p. 1001221. Available at: 10.1371/journal.pbio.1001221.

38. Martens, E.C., Chiang, H.C. and Gordon, J.I. (2008) ‘Mucosal Glycan Foraging Enhances Fitness and Transmission of a Saccharolytic Human Gut Bacterial Symbiont’, Cell Host and Microbe, 4(5), pp. 447–457. Available at: 10.1016/j.chom.2008.09.007.

39. Mukhopadhya, I. and Louis, P. (2025) ‘Gut microbiota-derived short-chain fatty acids and their role in human health and disease’, Nature Reviews Microbiology 2025, pp. 1–17. Available at: 10.1038/s41579-025-01183-w.

40. Ndeh, D. et al. (2017) ‘Complex pectin metabolism by gut bacteria reveals novel catalytic functions’, Nature, 544(7648), pp. 65–70. Available at: 10.1038/nature21725.

41. Ndeh, D.A. et al. (2025) ‘A Bacteroides thetaiotaomicron genetic locus encodes activities consistent with mucin O-glycoprotein processing and N-acetylgalactosamine metabolism’, Nature Communications 2025 16:1, 16(1), pp. 1–20. Available at: 10.1038/s41467-025-58660-2.

42. Neumann-Schaal, M. et al. (2015) ‘Time-resolved amino acid uptake of Clostridium difficile 630Δerm and concomitant fermentation product and toxin formation’, BMC Microbiology, 15(1), pp. 1–12. Available at: 10.1186/S12866-015-0614-2/FIGURES/6.

43. Ng, K.M. et al. (2013) ‘Microbiota-liberated host sugars facilitate post-antibiotic expansion of enteric pathogens’, Nature, 502(7469), pp. 96–99. Available at: 10.1038/nature12503.

44. Osbelt, L. et al. (2024) Klebsiella oxytoca inhibits Salmonella infection through multiple microbiota-context-dependent mechanisms, Nature Microbiology. Springer US. Available at: 10.1038/s41564-024-01710-0.

45. Passalacqua, K.D., Charbonneau, M.-E. and O’Riordan, M.X.D. (2016) ‘Bacterial Metabolism Shapes the Host–Pathogen Interface’, Microbiology Spectrum, 4(3). Available at: 10.1128/MICROBIOLSPEC.VMBF-0027-2015,.

46. Payling, L. et al. (2020) ‘The effects of carbohydrate structure on the composition and functionality of the human gut microbiota’, Trends in Food Science & Technology, 97, pp. 233–248. Available at: 10.1016/J.TIFS.2020.01.009.

47. Porter, N.T., Luis, A.S. and Martens, E.C. (2018) ‘Bacteroides thetaiotaomicron’, Trends in Microbiology, 26(11), pp. 966–967. Available at: 10.1016/j.tim.2018.08.005.

48. Rakoff-Nahoum, S., Coyne, M.J. and Comstock, L.E. (2014) ‘An ecological network of polysaccharide utilization among human intestinal symbionts’, Current Biology, 24(1), pp. 40–49. Available at: 10.1016/j.cub.2013.10.077.

49. Rivera-Chávez, F. and Bäumler, A.J. (2015) ‘The Pyromaniac Inside You: Salmonella Metabolism in the Host Gut’, Annual Review of Microbiology, 69(1), pp. 31–48. Available at: 10.1146/ANNUREV-MICRO-091014-104108,.

50. Rüttiger, A.-S. et al. (2025) ‘The global RNA-binding protein RbpB is a regulator of polysaccharide utilization in Bacteroides thetaiotaomicron’, Nature Communications, 16(1), p. 208. Available at: 10.1038/S41467-024-55383-8.

51. Ryan, D. et al. (2020) ‘A high-resolution transcriptome map identifies small RNA regulation of metabolism in the gut microbe Bacteroides thetaiotaomicron’, Nature Communications, 11(1). Available at: 10.1038/s41467-020-17348-5.

52. Ryan, D. et al. (2024) An expanded transcriptome atlas for Bacteroides thetaiotaomicron reveals a small RNA that modulates tetracycline sensitivity, Nature Microbiology. Springer US. Available at: 10.1038/s41564-024-01642-9.

53. Schulte, L.N. et al. (2011) ‘Analysis of the host microRNA response to Salmonella uncovers the control of major cytokines by the let-7 family’, EMBO Journal, 30(10), pp. 1977–1989. Available at: 10.1038/EMBOJ.2011.94,.

54. Schwalm, N.D., Townsend, G.E. and Groisman, E.A. (2016) ‘Multiple signals govern utilization of a polysaccharide in the gut bacterium bacteroides thetaiotaomicron’, mBio, 7(5). Available at: 10.1128/MBIO.01342-16,.

55. Shelton, C.D. et al. (2022) ‘Salmonella enterica serovar Typhimurium uses anaerobic respiration to overcome propionate-mediated colonization resistance’, Cell Reports, 38(1), p. 110180. Available at: 10.1016/j.celrep.2021.110180.

56. Smith, E.A. and Macfarlane, G.T. (1998) ‘Enumeration of amino acid fermenting bacteria in the human large intestine: effects of pH and starch on peptide metabolism and dissimilation of amino acids’, FEMS Microbiology Ecology, 25(4), pp. 355–368. Available at: 10.1111/J.1574-6941.1998.TB00487.X.

57. Sonnenburg, E.D. et al. (2010) ‘Specificity of polysaccharide use in intestinal bacteroides species determines diet-induced microbiota alterations’, Cell, 141(7), pp. 1241–1252. Available at: 10.1016/j.cell.2010.05.005.

58. Spiga, L. et al. (2017) ‘An Oxidative Central Metabolism Enables Salmonella to Utilize Microbiota-Derived Succinate’, Cell Host and Microbe, 22(3), pp. 291–301.e6. Available at: 10.1016/j.chom.2017.07.018.

59. Stecher, B. et al. (2007) ‘Salmonella enterica Serovar Typhimurium Exploits Inflammation to Compete with the Intestinal Microbiota’, PLOS Biology, 5(10), p. e244. Available at: 10.1371/JOURNAL.PBIO.0050244.

60. Townsend, G.E. et al. (2020) ‘A Master Regulator of Bacteroides thetaiotaomicron Gut Colonization Controls Carbohydrate Utilization and an Alternative Protein Synthesis Factor’, mBio, 11(1). Available at: 10.1128/MBIO.03221-19.

61. Tramontano, M. et al. (2018) ‘Nutritional preferences of human gut bacteria reveal their metabolic idiosyncrasies’, Nature Microbiology, 3(April), pp. 1–9. Available at: 10.1038/s41564-018-0123-9.

62. Varel, V.H. and Bryant, M.P. (1974) ‘Nutritional features of Bacteroides fragilis subsp. fragilis.’, Applied Microbiology, 28(2), pp. 251–257. Available at: 10.1128/AM.28.2.251-257.1974.

63. Vogel, J. (2020) ‘An RNA biology perspective on species-specific programmable RNA antibiotics’, Molecular Microbiology, 113(3), pp. 550–559. Available at: 10.1111/MMI.14476,.

64. Wardman, J.F. et al. (2022) ‘Carbohydrate-active enzymes (CAZymes) in the gut microbiome’, Nature Reviews Microbiology 2022 20:9, 20(9), pp. 542–556. Available at: 10.1038/s41579-022-00712-1.

65. Wende, M. et al. (2025) ‘Suppression of gut colonization by multidrug-resistant Escherichia coli clinical isolates through cooperative niche exclusion’, Nature communications, 16(1), p. 5426. Available at: 10.1038/S41467-025-61327-7.

66. Wexler, A.G. and Goodman, A.L. (2017) ‘An insider’s perspective: Bacteroides as a window into the microbiome.’, Nature microbiology, 2, p. 17026. Available at: 10.1038/nmicrobiol.2017.26.

67. Wexler, H.M. (2007) ‘Bacteroides: The good, the bad, and the nitty-gritty’, Clinical Microbiology Reviews, pp. 593–621. Available at: 10.1128/CMR.00008-07.

68. Whitaker, W.R., Shepherd, E.S. and Sonnenburg, J.L. (2017) ‘Tunable expression tools enable single-cell strain distinction in the gut microbiome’, Cell, 169(3), p. 538. Available at: 10.1016/J.CELL.2017.03.041.

69. Will, S.E. et al. (2019) ‘Day and Night: Metabolic Profiles and Evolutionary Relationships of Six Axenic Non-Marine Cyanobacteria’, Genome Biology and Evolution, 11(1), pp. 270–294. Available at: 10.1093/GBE/EVY275.

70. Winter, S.E. et al. (2013) ‘Host-derived nitrate boosts growth of E. coli in the inflamed gut’, Science, 339(6120), pp. 708–711. Available at: 10.1126/SCIENCE.1232467/SUPPL_FILE/WINTER.SM.PDF.

71. Wong, J.P.H. et al. (2023) ‘Bacteroides thetaiotaomicron metabolic activity decreases with polysaccharide molecular weight’.

72. Zampieri, M. et al. (2019) ‘Regulatory mechanisms underlying coordination of amino acid and glucose catabolism in Escherichia coli’, Nature Communications 2019 10:1, 10(1), pp. 1–13. Available at: 10.1038/s41467-019-11331-5.

73. Zhu, W. et al. (2020) ‘Xenosiderophore Utilization Promotes Bacteroides thetaiotaomicron Resilience during Colitis’, Cell Host & Microbe, 27(3), pp. 376–388.e8. Available at: 10.1016/j.chom.2020.01.010.

